# QTL Mapping on a Background of Variance Heterogeneity

**DOI:** 10.1101/276980

**Authors:** Robert W. Corty, William Valdar

## Abstract

Standard QTL mapping procedures seek to identify genetic loci affecting the phenotypic mean while assuming that all individuals have the same residual variance. But when the residual variance differs systematically between groups, perhaps due to a genetic or environmental factor, such standard procedures can falter: in testing for QTL associations, they attribute too much weight to observations that are noisy and too little to those that are precise, resulting in reduced power and and increased susceptibility to false positives. The negative effects of such “background variance heterogeneity” (BVH) on standard QTL mapping have received little attention until now, although the subject is closely related to work on the detection of variance-controlling genes. Here we use simulation to examine how BVH affects power and false positive rate for detecting QTL affecting the mean (mQTL), the variance (vQTL), or both (mvQTL). We compare linear regression for mQTL and Levene’s test for vQTL, with tests more recently developed, including tests based on the double generalized linear model (DGLM), which can model BVH explicitly. We show that, when used in conjunction with a suitable permutation procedure, the DGLM-based tests accurately control false positive rate and are more powerful than the other tests. We also find that some adverse effects of BVH can be mitigated by applying a rank inverse normal transform. We apply our novel approach, which we term “mean-variance QTL mapping”, to publicly available data on a mouse backcross and, after accommodating BVH driven by sire, detect a new mQTL for bodyweight.

## INTRODUCTION

A standard modeling assumption in quantitative trait locus (QTL) mapping is that all individuals, regardless of differences in their phenotypic mean, have the same residual variance. In reality, the residual variance (sometimes termed the environmental variance) between individuals can differ: the inherent noisiness of a phenotype can be affected by many factors, both extrinsic, such as environmental factors, and intrinsic, such as sex, or, more broadly, genetics. Environmental sources of residual variance heterogeneity have been well-documented, and include, for example, soil nitrogen and irrigation (Makumburage and Stapleton 2011), temperature (Shen *et al.* 2014), and even the age at which young birds begin to experience the environmental insults outside of the nest (Snell-Rood *et al.* 2015). Genetic sources of residual variance heterogeneity have attracted increasing interest, with multiple studies finding instances of the residual variance being heritable (Sorensen and Waagepetersen 2003; Hill and Mulder 2010; Sorensen *et al.* 2015; Gonzalez *et al.* 2016; Lin *et al.* 2016; Mitchell *et al.* 2016), and in some cases substantially attributable to allelic variation in individual genes (Paré *et al.* 2010; Wolc *et al.* 2012; Yang *et al.* 2012; Hulse and Cai 2013; Wang *et al.* 2014; Ayroles *et al.* 2015; Forsberg *et al.* 2015; Ivarsdottir *et al.* 2017).

The presence of residual variance heterogeneity, however, regardless of its source, can be problematic for analysis protocols that disregard it. Differences in residual variance between groups of individuals affect the precision of estimated means and, in turn, tests of significance or association (Cochran 1937; Yates and Cochran 1938). In the context of QTL mapping, ignoring such differences discards information that could be exploited to increase the power to detect QTL; and in the case of vQTL mapping specifically, it can covertly increase the false positive rate to well above the nominal level.

Specifically, the background presence of a major variance-controlling factor (*e.g.*, sex, housing, strain, a vQTL, etc.) implies that inferences about any other QTL effect (*e.g.*, that of a QTL elsewhere in the genome) occur against a backdrop of systematically heterogeneous residual variance. This “background variance heterogeneity” (BVH) acts to disrupt the natural observation weights: rather than every individual being subject to equal noise variance and therefore meriting equal weight, with BVH present some individuals’ phenotypes are inherently more (or less) noisy and so due less (or more) weight. Just as reweighting accordingly should lead to a more powerful analysis, assuming variance homogeneity— giving equal weight to subgroups of the data that are inherently noisier than average—risks overleveraging outliers and increasing the potential for both false negatives and false positives. This is likely to be true not only for studies detecting mQTL but also those detecting vQTL, which rely on the accurate attribution of residual noise.

Nonetheless, consideration of variance effects—whether as the target of inference or as a feature of the data to be accommodated— has thus far remained outside of routine genetic analysis. This could be in part because vQTL are sometimes considered of esoteric secondary interest, intrinsically controversial in their interpretation (Sun *et al.* 2013; Shen and Ronnegard 2013), or *a priori* too hard to detect (Visscher and Posthuma 2010). But it is also likely to be in part because standard protocols for finding and reporting vQTL are currently lacking, and because the advantages of modeling heterogeneous variance, even when targeting mQTL, remain under-appreciated and largely undemonstrated.

A number of statistical models and methods have been developed or adapted specifically to detect vQTL. These include: Levene’s test (Struchalin *et al.* 2010) and its generalizations (Soave *et al.* 2015; Soave and Sun 2017); the Fligner-Killeen test (Fraser and Schadt 2010); Bartlett’s test (Freund *et al.* 2013); and methods based on, or related to, the double generalized linear model (DGLM) and similar (Rönnegård and Valdar 2011; Cao *et al.* 2014; Dumitrascu *et al.* 2018). Tests have also been developed to detect genotype associations with arbitrary functions of the phenotype, for example higher moments, and these include a variant of the Komolgorov-Smirnov test (Aschard *et al.* 2013) and a semiparametric exponential tilt model (Hong *et al.* 2016). Of these, the ability to accommodate BVH of known source is limited to the DGLM of Rönnegård and Valdar (2011) (as well as a very recent Bayesian counterpart, described in Dumitrascu *et al.* 2018), which can include variance effects of arbitrary covariates as well as those belonging to the target (or foreground) QTL.

When the source of BVH is unknown, strategies to protect against it are less obvious. Since the threat manifests through sensitivity to distributional assumptions, possible remedies include side-stepping such assumptions via non-parametric approaches, *e.g.*, permutation testing, or reshaping the distribution prior to analysis through variable transformation. Both have been considered in the vQTL context, with permutation used in Hulse and Cai (2013) and Yang *et al.* (2012) and transformation in Rönnegård and Valdar (2011), Yang *et al.* (2012), Sun *et al.* (2013), and Shen and Carlborg (2013), but not specifically for controlling mQTL or vQTL false positives in the presence of BVH.

Here we examine the effect of modeled and unmodeled BVH on power and false positive rate when mapping QTL affecting the mean, the variance, or both. In doing so we:

1. Describe how the DGLM can be used develop a robust, straightforward procedure for routine mQTL and vQTL analysis, which we term “mean-variance QTL mapping”;
2. Compare alternative proposed methods for mQTL and vQTL analysis;
3. Show how accommodating BVH with the DGLM can improve power for detecting mQTL, vQTL, and mvQTL compared with other methods;
4. Show how sensitivity to model assumptions can be rescued by variable transformation and/or permutation; and
5. Demonstrate the discovery of a new QTL for mouse body-weight from an existing F2 intercross data resource (Leamy *et al.* 2000).

In two companion papers, we describe R package vqtl, which implements our procedure (Corty and Valdar 2018), and in Corty *et al.* (2018) apply it to two published QTL mapping experiments detecting a novel mQTL in one and a novel vQTL in the other. In particular, Corty *et al.* (2018) demonstrates a principle investigated here: that when an mQTL also has variance effects, those variance effects induce a type of proximal BVH, and modeling them explicitly therefore improves mQTL detection.

## STATISTICAL METHODS

This section reviews the tests and evaluation procedures that we studied through simulation. First, we describe eight statistical tests that can be used to model the effect of a single locus on phenotype mean and/or variance: the standard linear model, Levene’s test, Cao’s three tests, and three DGLM-based tests. We also describe four procedures for evaluating the statistical significance (*i.e.*, calculating p-values) of these tests—a standard asymptotic evaluation and three procedures that reasonably could be expected to provide protection against violations of model assumptions.

### Definitions

We start by defining three partially overlapping classes of QTL:

**mQTL:** a locus containing a genetic factor that causes heterogeneity of phenotype mean,
**vQTL:** a locus containing a genetic factor that causes heterogeneity of phenotype variance, and
**mvQTL:** a locus containing a genetic factor that causes heterogeneity of either phenotype mean, variance, or both — a generalization that includes the other two classes.

In addition, since we restrict our attention to QTL mapping methods that test genetic association with a phenotype one locus at a time, we distinguish two sources of variance effects:

**Foreground Variance Heterogeneity (FVH):** effects on the phenotype variance that arise from the locus under consideration (the focal locus);
**Background Variance Heterogeneity (BVH):** effects on the phenotype variance that arise from outside of the focal locus, *e.g.*, from another locus or an experimental covariate.

### Procedures to evaluate the significance of a single test

In comparing different statistical tests and their sensitivity to BVH, namely the effect of BVH on power and false positive rate (FPR), it is important to acknowledge that various measures could be taken to make significance testing procedures more robust to model misspecification in general and to BVH specifically. The significance testing methods considered here are frequentist, involving the calculation of a test statistic *T* on the observed data followed by an estimation of statistical significance based on a conception of *T*’s distribution under the null. However, BVH constitutes a departure of distributional assumptions, and in any rigorous applied statistical analysis when departures are expected it would be typical to consider protective measures such as, for example, transforming the response to make asymptotic assumptions more reasonable, or the use of computationally intensive procedures, such as those based on bootstrapping or permutation, to evaluate significance empirically.

Nominal significance (*i.e.,* the p-value for a single hypothesis test) is evaluated using four distinct procedures. The first two rely on asymptotics:

1. Standard: The test statistic *T* is computed on the observed data and compared with its asymptotic distribution under the null.
2. Rank-based inverse normal transform (RINT): As for standard, except observed phenotypes 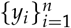 are first transformed to strict normality using the function RINT (*y_i_*) = Φ^−1^[(rank (*y_i_*) − ⅜) / (*n* + ¼)], where Φ is the normal c.d.f. and rank (*y_i_*) is gives the rank (from 1, …, *n*) (Beasley *et al*. 2009).

The second two determine significance empirically based on randomization: the test statistic *T* is recomputed as *T*^(*r*)^ under randomizations of the data *r* = 1, …, *R*, and the resulting set of statistics 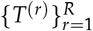 is used as the empirical distribution of *T* under the randomized null. Two alternative randomizations are considered:

3. Residperm: we generate a pseudo-null response 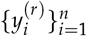 based on permuting the residuals of the fitted null model, (Freedman and Lane 1983; Good 2013), a process recently applied in the field of QTL mapping by Cao *et al*. (2014).
4. Locusperm: we leave the response intact, instead permuting the rows of the design matrix (or matrices) that differentiate(s) the null from alternative model.

### Procedure to evaluate genomewide significance

In the context of a genome scan, where many hypotheses are tested, we aim to control FPR genomewide through a family-wise error rate (FWER), the probability of making at least one false positive finding across the whole genome. This is done following the general approach of Churchill and Doerge (1994), which is closely related to the locusperm procedure described above, and which we refer to as genomeperm. Briefly, we perform an initial genome scan, recording test statistics 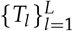 for all *L* loci. Then for each randomization *r* = 1, …, *R*, and for only the parts of the model that distinguish the null from the alternative model, the genomes are permuted among the individuals; the scan is then repeated to yield simulated null test statistics 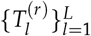 of which the maximum, 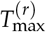, is recorded. The collection of 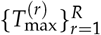 from all *R* such permutations is then used to fit a generalized extreme value distribution (GEV) (Dudbridge and Koeleman 2004; Valdar *et al*. 2006), and the quantiles of this are used to estimate FWER-adjusted p-values for each 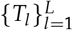.

### Standard linear model (SLM) for detecting mQTL

The standard model of quantitative trait mapping uses a linear regression based on the approximation of Haley and Knott (1992) and Martínez and Curnow (1992) to interval mapping of Lander and Botstein (1989). The effect of a given QTL on quantitative phenotype *y_i_* of individual *i* = 1, …, *n* is modeled as

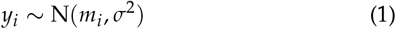

where *σ*^2^ is the residual variance and mi is a linear predictor for the mean, defined, in what we term the “full model”, as

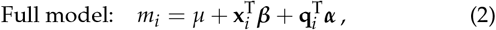

where *μ* is the intercept, *x_i_* is a vector of covariates with effects ***β***, and ***q***_*i*_ is a vector encoding the genetic state at the putative mQTL with corresponding mQTL effects ***α***. In the case considered here of biallelic loci arising from a cross of two founders, A and B, the genetic state vector ***q***_*i*_ = (*a_i_*, *d_i_*)^T^ is defined as follows: when genotype is known, for genotypes (AA,AB, BB), the additive dosage is *a_i_* = (0, 1, 2) and the dominance predictor is *d_i_* = (0, 1, 0); when genotype is available only as estimated probabilities *p*(AA), *p*(AB) and *p*(BB), following (Haley and Knott 1992; Martínez and Curnow 1992), we use the corresponding expectations, *a_i_* = 2*p*(AA) + *p*(AB) and *d_i_* = *p*(AB).

The test statistic for an mQTL is based on comparing the fit of the full model, acting as an alternative model, with that of a null that omits the locus effect, namely,

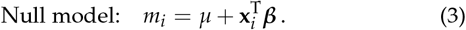

Since the regression in each case provides a maximum likelihood fit, the test statistic used here is likelihood ratio (LR) statistic, *T* = 2(*ℓ*_1_ – *ℓ*_0_), where *ℓ*_1_ and *ℓ*_0_ are the log-likelihoods under the alternative and the null respectively. For the biallelic model, the asymptotic test is the likelihood ratio test (LRT) whereby under the null, 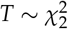. (Note: Alternative evaluation using the F-test is in general more precise but for our purposes provides equivalent results.)

The residperm approach to empirical significance evaluation of T proceeds as follows. We first fit the null model (Equation 3) to obtain predicted values 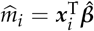 and estimated residuals *ε̂_i_* such that *y_i_* = *m̂_i_* + *ε̂_i_*. Then, for each randomization *r* = 1, …, *R*, we generate pseudo-null phenotypes 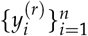 as

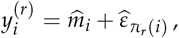

where if *π_r_* is a vector containing a random permutation of the indices *i* = 1, … *n*, then *π_r_* (i) is its *i*th element, mapping index *i* to its *r*th permuted version. The null and alternative models are then fitted to 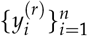 to yield 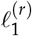 and 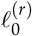, and hence *T*^(*r*)^.

In the locusperm approach to empirical significance, the response is unchanged but permutations are applied to the locus genotypes. For each randomization *r*, the full model *m_i_* is

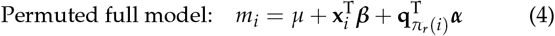

where *π_r_*(*i*) is as defined for residperm above. This full model fit yields 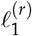, and then 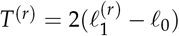. Note that 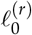 need not be recomputed after randomization because because only the rows of the design matrices that are unique to the alternative model are permuted and thus 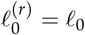.

### Levene’s Test (LV) for detecting vQTL

Levene’s test is a procedure for differences in variance between groups that can be used to detect vQTL. Suppose individuals are in G mutually exclusive groups *g* = 1, …, *G*. Let *g*[*i*] denote the group to which individual *i* belongs, denote *g*th group size as 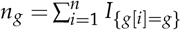, and *g*th group mean as 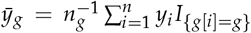. Then denote the *i*th absolute deviation as *z_i_* = |*y_i_* – *y*_*g*[*i*]_|, the group mean of these as 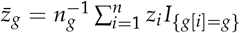 and overall mean 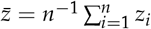. Levene’s W statistic is then

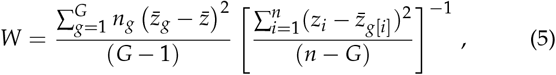

which under the null model of no variance effect follows the F distribution as *W* ~ *F*(*N* – *G*, *G* – 1) (Levene 1960). Note that replacing means of *y* with medians gives the related Brown-Forsythe test (Brown and Forsythe 1973), and replacing all instances of *z* with *y* in Equation 5 gives the ANOVA *F* statistic.

Levene’s test does not lend itself naturally to the residperm approach because it does not explicitly involve a null model to split the data into hat values and residuals. We therefore use the null model from the SLM (Equation 3) to approximate the residperm procedure with Levene’s test. To execute the locusperm procedure, for each randomization r, the group labels are permuted among the individuals, which is equivalent to replacing all instances of *g*[*i*] above with *g*[*π_r_*(*i*)], with *π_r_*(*i*) defined as above. A corresponding genomewide procedure, although not performed here, would ensure that each randomization *r* applies the same permutation *π_r_* across all loci.

### Cao’s Tests

Cao *et al*. (2014) elaborates the SLM to have a variance parameter that differs by genotype, *i.e.,*

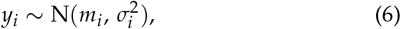

where *m_i_* is the linear predictor, 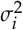 is the variance of the *i*th individual. These are defined in what we term the “full model” as

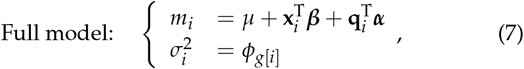

where *g*[*i*] indexes the genotype group to which *i* belongs, and 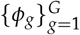 are the variances of the g = 1, …, *G* genotype groups. Thus an individual’s variance is entirely dictated by its genotype, and that genotype must be categorically known (or otherwise assigned). Cao *et al*. (2014) fits this model using a two-step, profile likelihood method, which in our applications we observe to be indistinguishable from full maximum likelihood (Figure S8).

Cao *et al.* (2014) describes tests for mQTL, vQTL and mvQTL based on comparing a full model against three different null models; we detail these tests below in our notation, denoting them respectively Cao_M_, Cao_V_, and Cao_MV_.

#### Cao_M_ test for detection of mQTL

The CaoM test involves an LRT between Cao’s full model and Cao’s no-mQTL model:

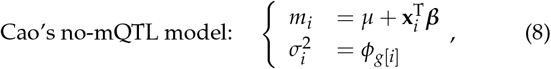

To execute the residperm procedure for Cao_M_, pseudo-null phenotypes are generated using *m̂_i_* and *ε̂_i_* from Cao’s no-mQTL model (Equation 8). The locusperm procedure respecifies the full model (Equation 7), leaving the variance model unchanged and specifying the mean predictor as 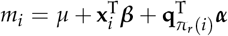.

#### Cao_V_ for detection of vQTL

The Cao_V_ test involves an LRT between Cao’s full model and Cao’s no-vQTL model:

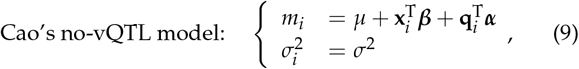

where the unsubscripted *σ*^2^ is a single, overall residual variance. This null model is identical to the alternative model in the SLM (Equation 2).

To execute the residperm procedure for Cao_V_, pseudo-null phenotypes are generating using *m̂_i_* and *ε̂_i_* from Cao’s no-mQTL model (Equation 9). The locusperm procedure respecifies the full model (Equation 7), leaving the mean sub-model unchanged and specifying the variance predictor as 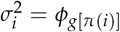.

#### Cao_MV_ for detection of generalized mvQTL

The Cao_MV_ test involves an LRT between Cao’s full model and Cao’s no-QTL model:

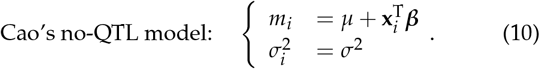

This null model is identical to the null model in the SLM (Equation 3).

To execute the residperm procedure for CaoMV, pseudo-null phenotypes are generated using *m̂_i_* and *ε̂_i_* from Cao’s no-QTL model (Equation 10). The locusperm procedure specifies the mean predictor as 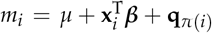 and the variance predictor as 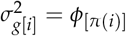.

### Double Generalized Linear Model (DGLM)

The DGLM models the phenotype *y_i_* via two linear predictors as

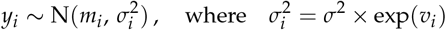

where *m_i_* predicts the phenotype mean and *υ_i_* predicts the extent to which the baseline residual variance *σ*^2^ is increased in individual *i*. In what we term the “DGLM full model”, these are specified as

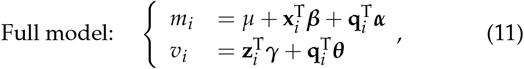

where *μ* is the intercept, **z**_*i*_ is a vector of covariates (which may beidentical to **x**_*i*_), *γ* is a vector of covariate effects on *υ_i_*, and ***θ*** is avector of locus effects on *υ_i_*.

As with Cao’s full model, the DGLM full model can be compared, in a likelihood ratio test, with various null models to test for mQTL, vQTL (Rönnegård and Valdar 2011), or mvQTL. A full maximum likelihood fitting procedure for the DGLM was provided by Smyth (1989).

#### DGLM_M_ for detecting mQTL

For detecting mQTL, we use an LRT of the DGLM full model in Equation 11 against the no-mQTL model:

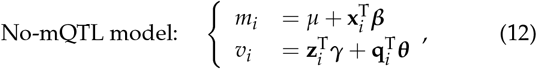

where the LR statistic has asymptotic distribution 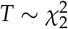.

To execute the residperm procedure for DGLMM, pseudo-null phenotypes are generated using *m̂_i_* and *ε̂_i_* from Equation 12. The locusperm procedure respecifies the mean predictor as 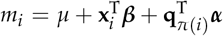 and does not modify the variance predictor.

#### DGLM_V_ for detecting vQTL

For detecting vQTL, we use an LRT of the DGLM full model in Equation 11 against the no-vQTL model:

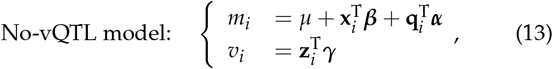

where the LR statistic has asymptotic distribution 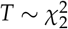.

To execute the residperm procedure for DGLMV, pseudo-null phenotypes are generated using *m̂_i_* and *ε̂_I_* from the Equation 13. The locusperm procedure does not modify the variance predictor and respecifies the mean predictor as 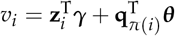.

#### DGLM_MV_ for detecting mvQTL

For detecting mvQTL, we use an LRT of the DGLM full model in Equation 11 against the no-QTL model:

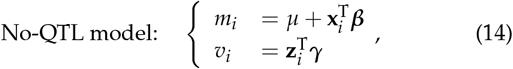

where the LR statistic has asymptotic distribution 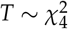.

To execute the residperm procedure for DGLM_MV_, pseudo-null phenotypes are generated using *m̂_i_* and *ε̂_i_* from the Equation 14. The locusperm procedure respecifies the mean predictor as 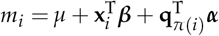 and the variance predictor as 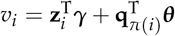.

## SIMULATION METHODS

The eight methods and four significance testing procedures described in the previous section, amounting to 32 test-procedure combinations in total, were compared by simulation. The simulations examined the performance of each combination, in terms of false and true positive rate, under eight distinct scenarios relating to the presence or absence of a QTL (and if present, then what type), and the presence or absence of BVH. We describe the general simulation setup below, followed by a detailed description of the eight scenarios and then describe the metrics by which performance was judged.

### Simulating locus and covariate

Each simulated experiment consisted of 300 individuals, where each individual was defined by one single-locus genotype, one covariate, and one phenotype.

The genotype for individual *i*, denoted by *q_i_*, was simulated according to a random process to mimic an F2 intercross:

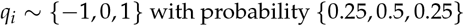

The covariate for individual *i*, denoted **z**_*i*_, was specified as a five-level categorical factor, with levels assigned to individuals as

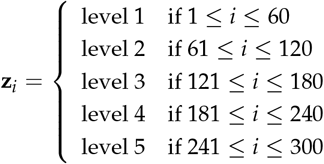

where **z**_*i*_ is an indicator vector such that, for example, **z***i* = (1, 0, 0, 0, 0) denotes membership of level 1. This covariate, which was fixed across simulations, was intended to mimic a generic, fixed aspect of experimental design in a typical QTL mapping study (for example, batch, technician, housing, etc.) that could plausibly influence the precision of the observations. When BVH is simulated, it is driven by this covariate.

### Scenarios

We conducted simulated experiments under eight different scenarios. These scenarios varied conceptually across two dimensions. First, we considered four types of locus:

1. null locus: The locus has no effect on phenotype.
2. pure mQTL: The locus has an additive effect on the phenotype mean.
3. pure vQTL: The locus has an additive effect on the log of the residual phenotype variance.
4. mixed mvQTL: The locus has both an additive effect on phenotype mean and an additive effect on the log of residual phenotype variance.

Then, we considered whether or not BVH was present, *i.e*.:

1. BVH absent: The covariate does not influence the residual variance of the phenotype.
2. BVH present: The covariate influences the residual variance of the phenotype (in addition to the locus, if a vQTL or mvQTL).

The resulting eight scenarios (*i.e*., all combinations) were realized *in silico* with three parameters: the effect of the locus on phenotype mean (*α*), the effect of the locus on phenotype variance (*θ*), and the effect of the covariate on phenotype variance (*γ*). Values assigned to these parameters are listed in Table 1. The rationale for selecting values of *α* and *θ* was as follows:

1. pure mQTL: The effect size of the pure mQTL was chosen so that it always explains 5% of the phenotype variance, which is consistent with smaller effect sizes typically sought and identified in QTL mapping experiments. Such an mQTL is detectable with approximately 70% power at a 5% false positive rate by the traditional mQTL test (the standard linear model) when 300 individuals are simulated, a typical population size for QTL mapping experiments.
2. pure vQTL: vQTL analysis is much less established, so the vQTL effect size was chosen to match the detectability of the mQTL. Thus, the vQTL effect size was defined such that the traditional vQTL test (Levene’s test) has 70% power at 5% FPR in a population of 300 individuals in the absence of BVH.
3. mixed mvQTL: The mvQTL effect sizes were chosen such that the mean and variance signals are equally detectable, and the aggregate signal is detectable by Cao_MV_ and DGLM_MV_ with 70% power at an FPR of 5% in a population of 300 individuals in the absence of BVH.

**Table 1.**
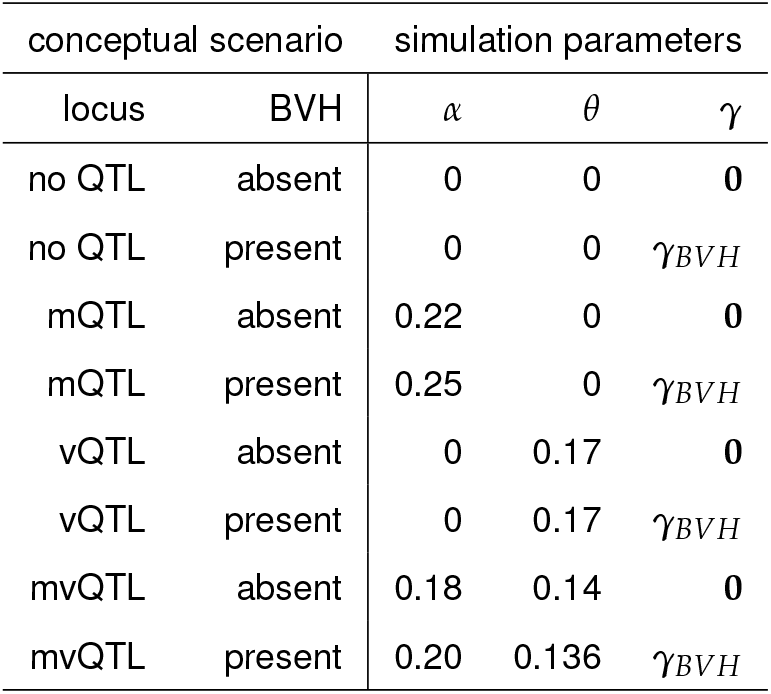
Eight scenarios were simulated, as determined by the values of three parameters. *α* indicates the additive effect of the locus on phenotype mean, *θ* the additive effect of the locus on phenotype variance, and *γ* the effect of the covariate on phenotype variance. The two possible values of *γ* are **0** = [0, 0, 0, 0, 0] and *γ_BVH_* = [–0.4,–0.2, 0, 0.2, 0.4].

The values of *γ* used for simulating BVH were **0** = [0, 0, 0, 0, 0] and *γ_BVH_* = [–0.4,–0.2, 0, 0.2, 0.4]. The former chosen to ensure constant residual variance for simulations where BVH is absent; the latter to mirror the extent of BVH we noted in experimental data, while having a concise expression as equally spaced effects centered at zero. In null locus and mQTL simulations, *γ_BVH_* results in group-wise standard deviations of approximately [0.67, 0.82, 1.00, 1.22, 1.49]. In vQTL and mvQTL simulations, *γ_BVH_* and *θ* combine additively on the log standard deviation scale and result in fifteen unique variances as detailed in the **Supplementary Materials.**

### Phenotype simulation

For each of the eight scenarios, we conducted 10, 000 simulated experiment. For scenario *s*, the phenotype for individual *i*, denoted *y_i_*, was simulated from a normal distribution based on the genotype and covariate (*q_i_* and *x_i_*) and the scenario parameters (*α_s_*, *θ_s_*, and *γ_s_*) as:

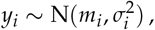

where *m_i_* = *q_i_α_s_*, and

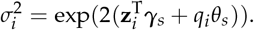

(Further details in **Supplementary Materials.**)

### Testing significance

To each simulated experiment, eight tests were applied, and four procedures were used to assess the statistical significance of each test, for a total of 32 test-procedures.

The eight tests comprise three tests for detecting mQTL: SLM, CaoM, and DGLMM; three for detecting vQTL: Levene’s test, CaoV, DGLMV; and two for detecting mvQTL: CaoMV and DGLMMV. These tests are detailed in the **Statistical Methods** and summarized in Table 2.

**Table 2.** The eight tests that were evaluated in the simulation studies. FVH: foreground variance heterogeneity (*i.e.*, variance effects at the QTL). BVH: background variance heterogeneity.

| Category | Test | Description |
| --- | --- | --- |
| mQTL | SLM | Conventional test of mean differences; allows neither FVH nor BVH |
| mQTL | Cao<sub>M</sub> | Allows FVH, but not BVH |
| mQTL | DGLM<sub>M</sub> | Allows FVH and BVH |
| vQTL | Levene's test | Conventional test of variance differences; detects FVH, does not allow BVH |
| vQTL | Cao<sub>V</sub> | Detects FVH, does not allow BVH |
| vQTL | DGLM<sub>V</sub> | Detects FVH, allows BVH |
| mvQTL | Cao<sub>MV</sub> | Detects FVH, does not allow BVH |
| mvQTL | DGLM<sub>MV</sub> | Detects FVH and allows BVH |

The four procedures for evaluating the statistical significance of results were: standard, RINT, residperm, and locusperm, as described in the **Statistical Methods**. The RINT procedure was selected because it returns any phenotype distribution, no matter how exotic, to a standard normal distribution. The fact that it is commonly used in genetics research demands that its properties, and its effects on QTL mapping, be better understood. The residperm was selected because it was recently proposed for use in mQTL, vQTL, and mvQTL mapping studies (Cao *et al.* 2014). The locusperm procedure was developed in response to suspected shortcomings of the above robustifying procedures.

### Evaluation of tests and procedures

Tests and procedures for assessing statistical significance were evaluated based on their empirical false positive rate (FPR) and power at a nominal FPR of 0.05. The empirical FPR of a given test-procedure combination in a given scenario is simply the fraction of null simulations (where the phenotype was simulated with no dependence on genotype) that resulted in *p* < 0.05. Similarly, the empirical power was computed as the fraction of non-null simulations that resulted in *p* < 0.05. These quantities are naturally considered as estimates of a binomial proportion, so their standard errors were calculated by the method of Clopper and Pearson (1934).

These evaluations focus only on the cutoff of *p* = 0.05. We considered all possible cutoffs with QQ plots and ROC plots. These evaluations examine the empirical FPR as a function of nominal FPR and the empirical power as a function of empirical FPR, respectively. They enrich the understanding of the spectrum of trade-offs that each test makes available, but do not meaningfully change the overall interpretation of the results, so we relegate them to the **Supplementary Materials**.

## DATA AND SOFTWARE

### Leamy *et al.* summary of original study

Leamy *et al.* (2000)
 backcrossed mice from strain CAST/Ei, a small, lean strain, into mouse strain M16i, a large, obese strain. Nine F1 males were bred with 54 M16i females to produce a total of 421 offspring (208 female, 213 male), which were genotyped at 92 microsatellite markers across the 19 autosomes and phenotyped for body composition and morphometric traits. We retrieved all available data on this cross, which included marker genotypes, covariates, and eight phenotypes (body weight at five ages, liver weight, subcutaneous fat pad thickness, and gonadal fat pad thickness), from the Mouse Phenome Database (Grubb *et al.* 2014), and estimated genotype probabilities at 2cM intervals across the genome using the hidden Markov model in R/qtl (Broman *et al.* 2003).

This mapping population has been studied for association with several phenotypes: asymmetry of mandible geometry (Leamy *et al.* 2000), limb bone length (Leamy *et al.* 2002; Wolf *et al.* 2006), organ weight (Leamy *et al.* 2002; Wolf *et al.* 2006; Yi *et al.* 2006), fat pad thickness (Yi *et al.* 2005, 2006, 2007), and body weight (Yi *et al.* 2006). The most relevant prior study to this reanalysis, Yi *et al.* (2006), used standard methods to identify QTL for body weight at three weeks on chromosomes 1 and 18. However, we were not able to reproduce this result, despite following their analysis as described.

### Availability of data and software

Analyses were conducted in the R statistical programming language (R Core Team 2017). The simulation studies used the implementation of the standard linear model from package stats, Levene’s test from car, Cao’s tests as published in Cao *et al.* (2014) and the DGLM tests in package dglm. Files S1, S2, and S3 contain the R scripts necessary to replicate the simulation studies and their analysis, relying on the plotROC package to make ROC plots (Sachs and Others 2017). File S4 contains the data from Leamy *et al.* (2000) that was reanalyzed. File S5 contains the attempted replication of the original analysis (Yi *et al.* 2006) and file S6 contains the new analysis, using package vqtl.

The reanalyzed dataset is available on the Mouse Phenome Database (Grubb *et al.* 2014) with persistent identifier MPD:206. The entire project, including data and all analysis scripts, is available as a public, static Zenodo repository with DOI:10.5281/zenodo.1455184.

## RESULTS

### Simulation study on single locus testing

Simulations were performed to examine the ability of the eight tests listed in Table 2 to detect nonzero effects belonging to their target QTL types (mQTL, vQTL, mvQTL), and to control the number of false positives when no such QTL effects were present. Simulations were conducted in the presence and absence of background variance heterogeneity (BVH), and for each test, with p-values calculated by each of the four significance assessment procedures (standard, RINT, residperm, locusperm). The full combination of settings is listed in Table 3, which also lists results pertaining to a nominal FPR of 0.05, and described in more detail in **Simulation Methods** section.

**Table 3.** Empirical positive rates of all tests under all significance assessment procedures in all scenarios based on 10,000 simulations, 1,000 permutations each to estimate empirical null distributions (residperm and locusperm), and a nominal false positive rate (FPR) of *α* = 0.05. Entries in column 1 and 5 through all rows, columns 3 and 7 in the top third, and columns 2 and 6 in the middle third represent empirical FPR. Where the empirical FPR is within one standard error of the nominal FPR of 0.05, it is written in normal font. Where it is overly conservative, it is underlined. Where it is anti-conservative, it is in boldface. The entries in the rest of the table represent power. Given the sample size of 10,000, the standard error for the values in this table are all between 0.005 and 0.01. Generally, values near 0.05 have a standard error near 0.005 and values near 0.5 have a standard error near 0.01. All standard errors are listed in table Table S2 and plotted visually in Figure 1, Figure 2, Figure 3, and Figure 4.

| test | procedure | BVH absent |  |  |  | BVH present |  |  |  |
| --- | --- | --- | --- | --- | --- | --- | --- | --- | --- |
|  |  | null | mQTL | vQTL | mvQTL | null | mQTL | vQTL | mvQTL |
| SLM | standard | 0.052 | 0.717 | 0.054 | 0.502 | 0.052 | 0.706 | 0.052 | 0.506 |
|  | RINT | 0.051 | 0.712 | 0.051 | 0.486 | 0.053 | 0.719 | 0.049 | 0.512 |
|  | residperm | 0.050 | 0.710 | 0.052 | 0.494 | 0.050 | 0.700 | 0.052 | 0.498 |
|  | locusperm | 0.049 | 0.709 | 0.052 | 0.497 | 0.051 | 0.699 | 0.051 | 0.499 |
| Cao<sub>M</sub> | standard | 0.053 | 0.717 | 0.051 | 0.510 | 0.054 | 0.702 | 0.049 | 0.520 |
|  | RINT | 0.052 | 0.713 | 0.048 | 0.496 | 0.054 | 0.717 | 0.049 | 0.529 |
|  | residperm | 0.051 | 0.709 | 0.048 | 0.501 | 0.051 | 0.698 | 0.048 | 0.509 |
|  | locusperm | 0.049 | 0.704 | 0.047 | 0.496 | 0.050 | 0.694 | 0.046 | 0.506 |
| DGLM<sub>M</sub> | standard | **0.057** | 0.714 | 0.055 | 0.510 | **0.059** | 0.836 | 0.057 | 0.654 |
|  | RINT | **0.055** | 0.713 | 0.055 | 0.499 | **0.057** | 0.834 | 0.056 | 0.648 |
|  | residperm | 0.049 | 0.693 | 0.048 | 0.490 | 0.052 | 0.822 | 0.050 | 0.631 |
|  | locusperm | 0.049 | 0.691 | 0.047 | 0.484 | 0.050 | 0.818 | 0.048 | 0.628 |
| Levene's test | standard | 0.045 | 0.048 | 0.655 | 0.452 | 0.050 | 0.045 | 0.566 | 0.388 |
|  | RINT | 0.046 | 0.041 | 0.638 | 0.416 | 0.048 | 0.041 | 0.539 | 0.340 |
|  | residperm | 0.049 | 0.052 | 0.664 | 0.461 | 0.052 | 0.048 | 0.574 | 0.397 |
|  | locusperm | 0.047 | 0.051 | 0.665 | 0.461 | 0.052 | 0.049 | 0.576 | 0.399 |
| Cao<sub>v</sub> | standard | **0.054** | 0.053 | 0.738 | 0.536 | **0.135** | 0.131 | 0.736 | 0.568 |
|  | RINT | <u>0.045</u> | 0.041 | 0.692 | 0.457 | 0.046 | 0.047 | 0.552 | 0.366 |
|  | residperm | 0.050 | 0.049 | 0.721 | 0.510 | 0.053 | 0.050 | 0.564 | 0.390 |
|  | locusperm | 0.049 | 0.047 | 0.717 | 0.505 | 0.051 | 0.047 | 0.561 | 0.384 |
| DGLM<sub>v</sub> | standard | **0.054** | 0.053 | 0.721 | 0.520 | **0.058** | 0.050 | 0.732 | 0.527 |
|  | RINT | <u>0.045</u> | 0.041 | 0.673 | 0.444 | <u>0.023</u> | 0.022 | 0.564 | 0.346 |
|  | residperm | 0.049 | 0.049 | 0.699 | 0.496 | <u>0.020</u> | 0.018 | 0.537 | 0.331 |
|  | locusperm | 0.048 | 0.048 | 0.698 | 0.490 | 0.052 | 0.046 | 0.708 | 0.502 |
| Cao<sub>MV</sub> | standard | 0.053 | 0.607 | 0.633 | 0.742 | **0.113** | 0.642 | 0.643 | 0.749 |
|  | RINT | 0.046 | 0.595 | 0.567 | 0.698 | 0.049 | 0.603 | 0.434 | 0.650 |
|  | residperm | 0.049 | 0.597 | 0.606 | 0.726 | 0.054 | 0.516 | 0.503 | 0.632 |
|  | locusperm | 0.049 | 0.596 | 0.610 | 0.729 | 0.053 | 0.517 | 0.503 | 0.634 |
| DGLM<sub>MV</sub> | standard | **0.056** | 0.606 | 0.621 | 0.734 | **0.060** | 0.746 | 0.631 | 0.811 |
|  | RINT | 0.050 | 0.597 | 0.559 | 0.694 | <u>0.038</u> | 0.724 | 0.440 | 0.732 |
|  | residperm | 0.050 | 0.589 | 0.587 | 0.711 | <u>0.025</u> | 0.631 | 0.466 | 0.694 |
|  | locusperm | 0.050 | 0.586 | 0.585 | 0.713 | 0.054 | 0.735 | 0.596 | 0.790 |

### Testing for mQTL with BVH absent: SLM and Cao_M_ outperform DGLM_M_

In the absence of BVH, SLM and Cao_M_ accurately control FPR under all significance assessment procedures (Figure 1 and Table 3). DGLMM was slightly anti-conservative under the standard and RINT procedures with FPR = 0.057 and 0.055, respectively. With either permutation procedure used to assess significance, however, DGLMM accurately controlled FPR.

**Figure 1.**
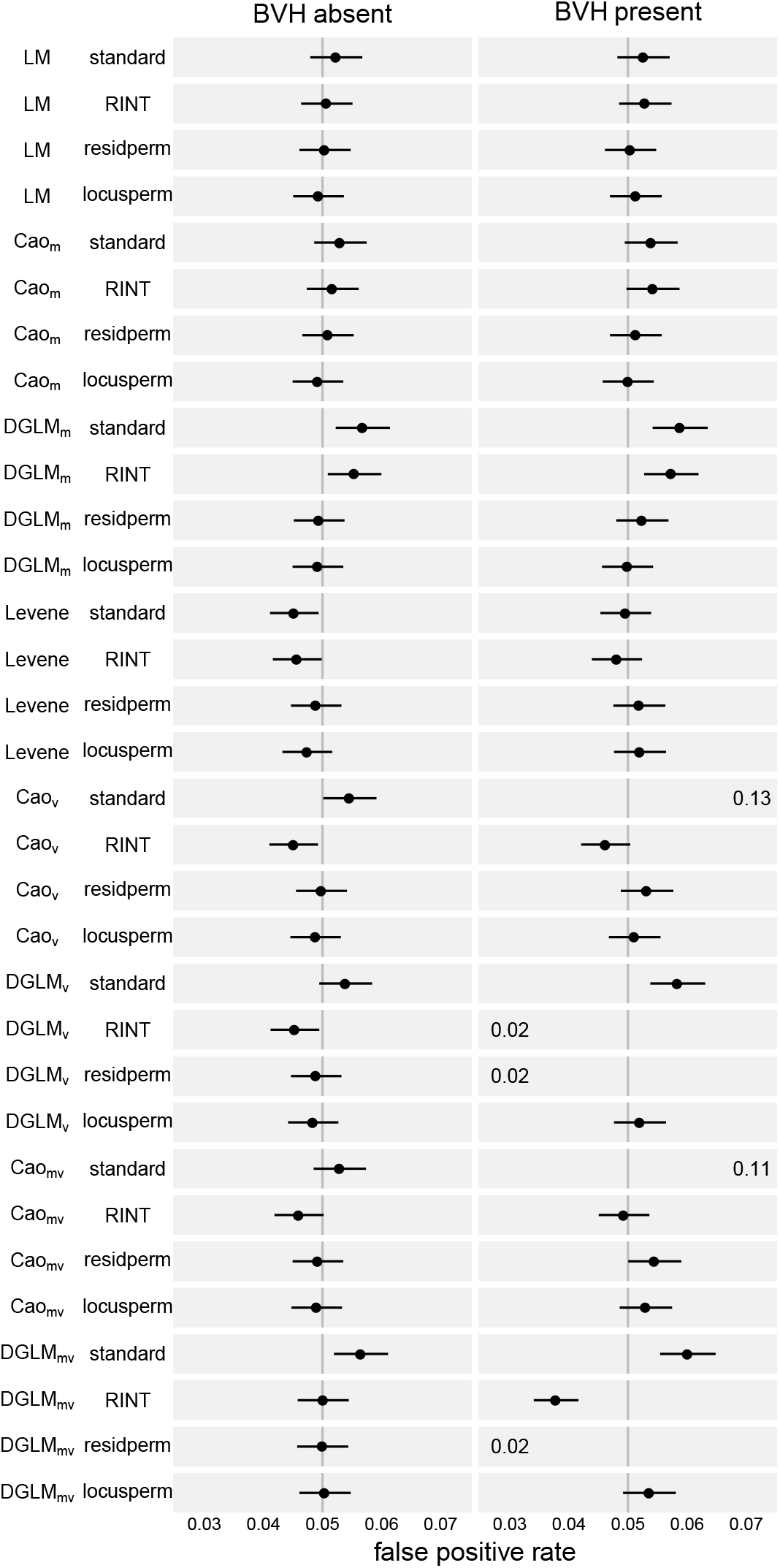
Empirical false positive rate (FPR) of all tests and significance assessment procedures at a nominal FPR of 0.05, as assessed through simulation of non-associated loci and phenotypes both with and without BVH. Dot indicates point estimate and line indicates 95% confidence interval. The vertical line indicates the ideal empirical FPR of 0.05. Some test-procedure combinations led to FPR outside the plotted range. In such cases the FPR is written on the left edge of the plotting area if the value was too low to plot, and the right edge if it was too high. An un-zoomed version of this plot is available in Figure S7.

SLM and Cao_M_ had indistinguishable power in the detection of mQTL under all significance assessment procedures (Figure 2). DGLM_M_, however, had equal power to those tests only under the standard and RINT procedures, which have inflated FPR. Under the permutation-based procedures, DGLM_M_ was less powerful than the other test-procedures.

**Figure 2.**
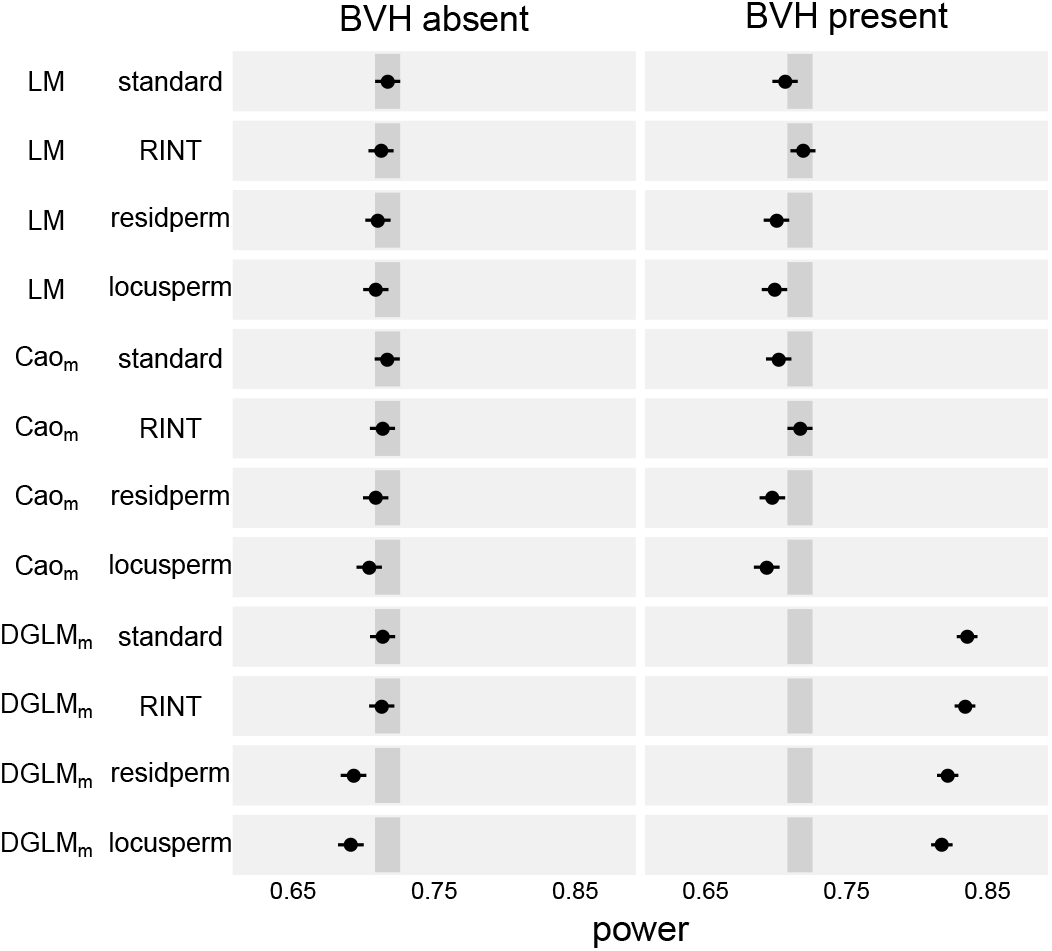
Empirical power of mQTL tests to detect mQTL under four significance assessment procedures. Dot indicates point estimate and line indicates 95% confidence interval. Darker gray rectangle indicates the confidence band for the power of SLM with the standard significance assessment procedure, the standard against which all other test-procedures are compared..

These results reflect the reality that, when a simple model is exactly true, a more elaborate model tends to be less powerful. Additionally, they highlight the capability of the permutation-based procedures to accurately control FPR even when the standard and RINT procedures fail to do so (as in the case of DGLM_M_).

#### Testing for mQTL with BVH present: DGLM_M_ dominates

SLM and CaoM accurately controlled FPR under all four procedures to assess statistical significance (Figure 1). As in the absence of BVH, DGLMM exhibited a slightly inflated FPR under the standard and RINT procedures (0.059 and 0.057, respectively), but accurately controlled FPR under the permutation-based procedures (Table 3).

Under all four procedures, DGLM_M_ was more powerful than SLM and CaoM (Figure 2). The two procedures under which DGLM_M_ accurately controlled FPR had power of 0.822 and 0.818, greatly exceeding the power of Cao_M_ and SLM, which were in the range [0.694, 0.719] (Table 3).

Based on the results of these simulations, DGLM_M_-residperm and DGLMM-locusperm are the recommended test-procedure combinations for mQTL testing in the presence of BVH.

For each mQTL test-procedure combination, the AUC (Table S1), standard error of the positive rate at *α* = 0.05 (Table S2), QQ plots illustrating the empirical FPR at each nominal FPR level (Figure S4), and ROC curves illustrating the spectrum of trade-offs between available FPR and power (Figure S1) are provided in the **Supplementary Materials**.

#### Testing for vQTL with BVH absent: Cao_V_ and DGLM_V_ outperform Levene’s test

In the absence of BVH, all vQTL tests had nearly-accurate FPR control (Figure 1). All tests had FPR within one standard error of 0.05 under both empirical significance assessment procedures (Table 3 and Table S2). But under either asymptotic procedure, Levene’s test was slightly conservative. And Cao_V_ and DGLM_V_ were both slightly anti-conservative under the standard procedure and conservative under the RINT procedure.

Despite the variation in FPR control among the test-procedure combinations, CaoV and DGLM_V_ had more power to detect vQTL than Levene’s test under all procedures. Specifically, under the well-calibrated (empirical) procedures, Cao_V_ and DGLM_V_ had power in the range [0.698, 0.721], while under those same well-calibrated (empirical) procedures, Levene’s test had power in the range [0.664, 0.665] (Table 3).

Thus, in the specific situations simulated here, the empirical procedures of Cao_V_ and DGLM_V_ are the preferred vQTL tests in the absence of BVH. The additional power of Cao_V_ and DGLM_V_ relative to Levene’s test is consistent with the fact that they make strong parametric assumptions that are exactly true in these simulations and Levene’s test does not.

#### Testing for vQTL with BVH present: DGLM_V_ outperforms Levene’s test and Cao_V_

In the presence of BVH, there were three test-procedure combinations with major departures from accurate FPR control (Figure 3). Cao_V_ under the standard procedure was drastically anti-conservative with FPR of 0.135 (Table 3). DGLM_V_ under both the RINT and residperm procedures was drastically conservative with FPR of 0.023 and 0.020, respectively. Additionally, DGLM_V_ under the standard procedure was moderately anti-conservative with FPR of 0.058. The remaining test-procedure combinations accurately controlled FPR, namely Levene’s test under all procedures, Cao_V_ under the RINT, residperm, and locusperm procedures, and DGLM_V_ under the locusperm procedure.

**Figure 3.**
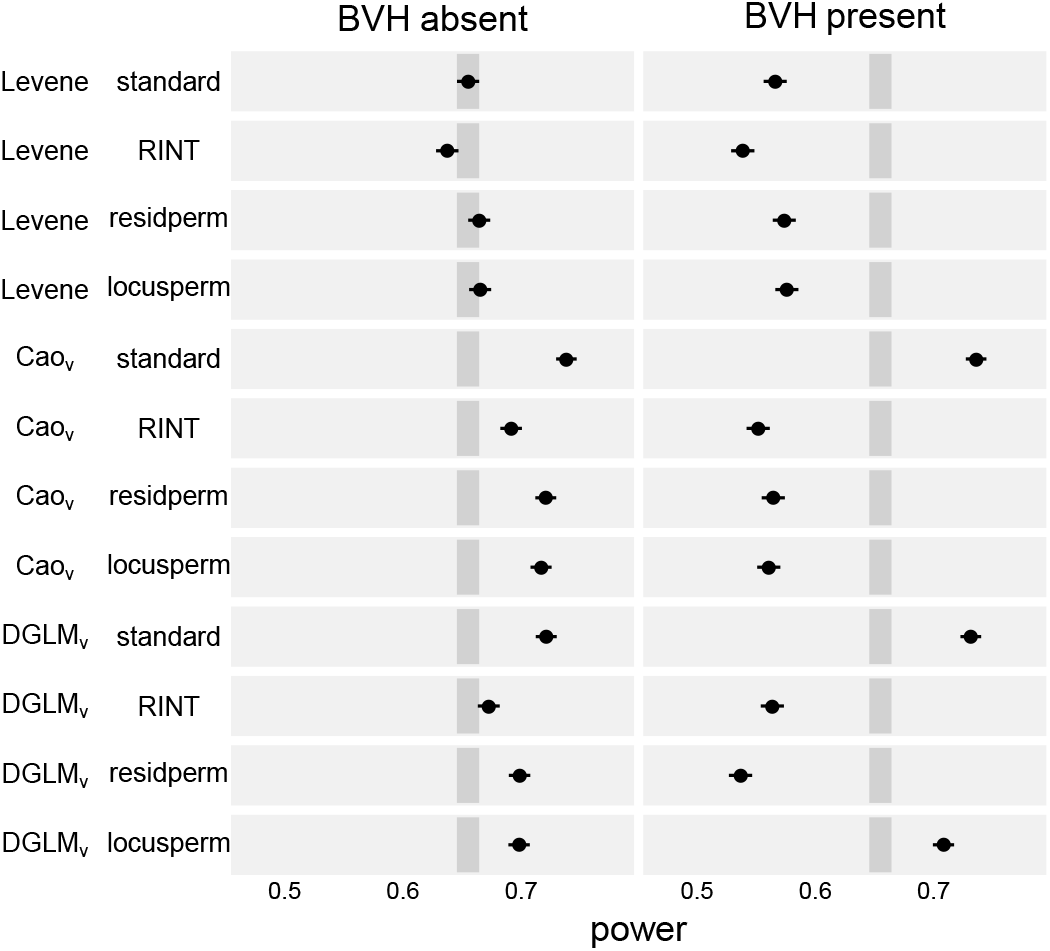
Empirical power of vQTL tests to detect vQTL under four significance assessment procedures. Dot indicates point estimate and line indicates 95% confidence interval. Darker gray rectangle indicates the confidence band for the power of Levene’s test with the standard significance assessment procedure, the standard against which the other test-procedures are compared.

Of the tests that accurately controlled FPR, DGLM_V_ under the locusperm procedure was uniquely powerful, with power of 0.708, while the others had power in the range [0.539, 0.576] (Figure 3 and Table 3).

Direct interpretation of these results might lead a geneticist to consider the trade-off between DGLM_V_-standard and DGLM_V_-locusperm. DGLM_V_-locusperm requires considerable computational effort and serves only to reduce the FPR from a modestly-inflated level of 0.058 to accurate control at 0.052. Application of DGLMV-standard, however comes with a dangerous caveat. If there were some additional, unknown (and therefore unmodeled) BVH-driving factor, DGLMV-standard should be expected to have drastically anti-conservative behavior, similar to Cao_V_-standard in the simulations described here in which BVH was present. The locusperm procedure, in contrast, ensures accurate FPR control whether all BVH-driving factors are modeled (as in DGLM_V_) or not (as in Cao_V_). Therefore, DGLMV-locusperm is the recommended test for vQTL mapping in the presence of BVH.

For each vQTL test-procedure combination, the AUC (Table S1), standard error of the positive rate at *α* = 0.05 (Table S2), QQ plots illustrating the empirical FPR at each nominal FPR level (Figure S5), and ROC curves illustrating the spectrum of trade-offs between available FPR and power (Figure S2) are provided in the **Supplementary Materials**.

#### Testing mvQTL with BVH absent: Cao_MV_ and DGLM_MV_ similar

Continuing the pattern from the vQTL tests, in the absence of BVH most mvQTL tests accurately control FPR (Figure 1). The exceptions are similar to the vQTL tests as well, with CaoMV-RINT slightly conservative and DGLMMV-standard slightly anti-conservative (Table 3).

The standard version of Cao_MV_ and DGLM_MV_ were similarly powerful (Figure 4), both exceeding the power of the other mvQTL test-procedures.

**Figure 4.**
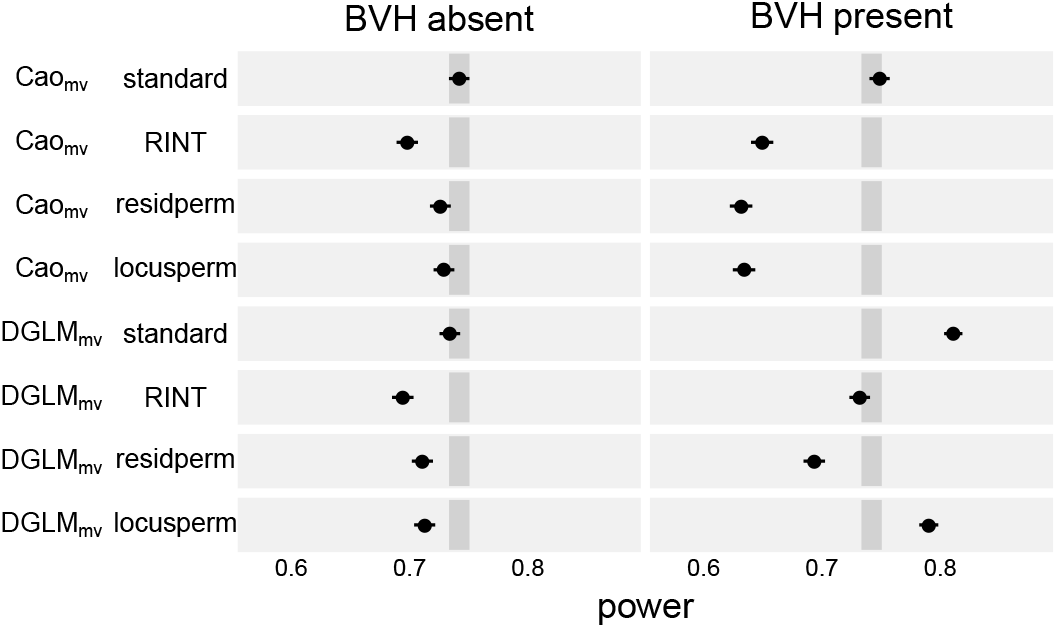
Empirical power of mvQTL tests to detect mvQTL under four significance assessment procedures. Dot indicates point estimate and line indicates 95% confidence interval. Darker gray rectangle indicates the confidence band for the power of CaoMV with the standard significance assessment procedure, the standard against which the other test-procedures are compared.

#### Testing mvQTL with BVH present: DGLM_MV_ dominates Cao_MV_

In the presence of BVH, Cao_MV_ accurately controlled FPR with the RINT, residperm, and locusperm procedures, while DGLM_MV_ did so only under the locusperm procedure (Figure 1).

Of the test-procedure combinations that accurately controlled FPR, DGLM_MV_-locusperm was the most powerful with power of 0.790 as compared to the others in the range [0.632, 0.650].

As with vQTL tests, the DGLM_MV_-standard appears to be a compelling balance between computational effort and good statistical properties, but it is expected to have drastically inflated FPR in the presence of any unmodeled BVH-driving factor, similar to Cao_MV_-standard. Therefore, DGLM_MV_-locusperm is the recommended test-procedure for mvQTL testing.

For each mvQTL test-procedure combination, the AUC (Table S1), standard error of the positive rate at *α* = 0.05 (Table S2), QQ plots illustrating the empirical FPR at each nominal FPR level (Figure S6), and ROC curves illustrating the spectrum of trade-offs between available FPR and power (Figure S3) are provided in the **Supplementary Materials**.

#### In the presence of BVH, the rank-based inverse normal transformation fails to correct anti-conservative behavior of DGLM_M_ and overcorrects that of DGLM_V_ and DGLM_MV_

A consistent feature of the simulations involving detection of variance effects, whether vQTL or mvQTL, is that FPR control and power is affected, for better or worse, by applying the RINT to the response.

In the presence of BVH, DGLMM under the standard procedure was anti-conservative (FPR = 0.059 at *α* = 0.05). The RINT procedure had little efficacy in returning this test to accurate FPR control (FPR = 0.057).

In the case of vQTL detection in the presence of BVH, Cao_V_ under the standard procedure had a drastically inflated FPR (0.135) and the RINT procedure slightly over-corrected it (FPR = 0.046). Similarly, the RINT procedure disrupted DGLM_V_, which was modestly anti-conservative under the standard procedure, causing overly conservative behavior (FPR = 0.023).

As always, in the presence of BVH, the mvQTL tests exhibited a mixture of the patterns observed in mQTL tests and vQTL tests. Both Cao_MV_ and DGLM_MV_ were anti-conservative under the standard procedure, illustrating their relations to CaoV and DGLMM respectively. In the case of Cao_MV_, the RINT procedure corrected the FPR, but in in the case of DGLM_MV_, it resulted in an over-correction into the realm of over conservatism (FPR = 0.049 and 0.038 respectively).

In summary, the RINT procedure is unhelpful in the context of the DGLMM: it does not repair the modest FPR inflation that is present under the standard procedure. But, in the context of vQTL testing with BVH, it has one useful and important property: pre-processing the phenotype with the RINT, leads to vQTL tests that are conservative rather than anti-conservative, decreasing the probability of false positives at the expense of false negatives.

### Genomewide reanalysis of bodyweight in Leamy et al. back-cross

To understand the impact of BVH on mean and variance QTL mapping in real data, we applied both traditional QTL mapping, using SLM, and mean-variance QTL mapping, using Cao’s tests and the DGLM, to body weight at three weeks in the mouse backcross dataset of Leamy *et al.* (2000).

#### Analysis with traditional QTL mapping identifies no QTL

We first used a traditional, linear modeling-based QTL analysis, with sex and father as additive covariates and genomewide significance based on 1000 genome permutations (Churchill and Doerge 1994). Although sex was found not to be a statistically significant predictor of body weight (*p* = 0.093 by the likelihood ratio test with 1 degree of freedom), it was included in the mapping model because, based on the known importance of sex in determining body weight, any QTL that could only be identified in the absence of modeling sex effects would be highly questionable. Father was found to be a significant predictor of body weight in the baseline fitting of the SLM (*p* = 9.6 × 10^−5^ by the likelihood ratio test with 8 degrees of freedom) and therefore was included in the mapping model.

No associations rose above the threshold that controls family-wise error rate to 5% (Figure 5, green line). One region on the distal part of chromosome 11 could be considered “suggestive” with FWER-adjusted *p* ≈ 0.17.

**Figure 5.**
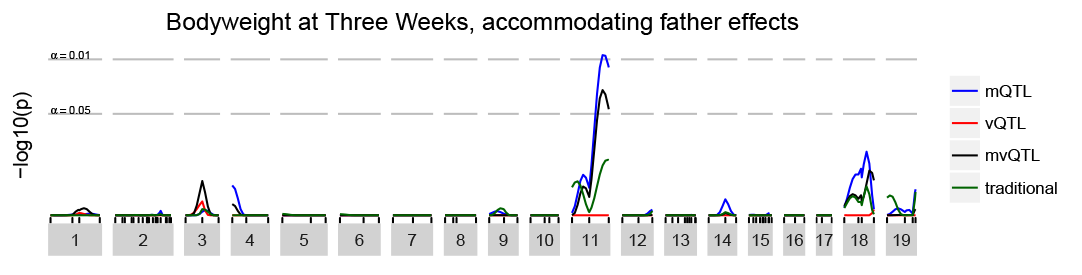
FWER-controlling association statistic at each genomic locus for body weight at three weeks. The linear model (green, “traditional”) does not detect any statistically-significant associations. The mQTL test takes into account the heterogeneity of both mean and variance due to which F1 male fathered each mouse in the mapping population and detects one mQTL on chromosome 11.

To test the sensitivity of the results to the inclusion/exclusion of covariates, the analysis was repeated without sex as a covariate, without father as a covariate, and with no covariates. No QTL were identified in any of these sensitivity analyses.

#### Analysis with Cao’s tests identifies no QTL

The same phenotype was analyzed with Cao’s tests, again including sex and father as mean covariates, and using the genome permutation procedures described in **Statistical Methods** were used to control FWER. No statistically significant mQTL, vQTL, nor mvQTL were identified (Figure S10).

#### Analysis with DGLM-based tests identifies an mQTL

The same phenotype was analyzed with the DGLM-based tests. In a baseline fitting of the DGLM, sex was found not to be a statistically significant predictor of mean or residual variance (mean effect *p* = 0.18, variance effect *p* = 0.22, and joint *p* = 0.19 by the LRT with 1, 1, and 2 d.f.). But father was found to be a statistically significant predictor of both mean and variance (mean effect *p* = 2.0 × 10^−7^, variance effect *p* = 1.8 × 10^−11^, and *p* = 4.8 × 10^−14^ by the LRT with 8, 8, and 16 d.f.). Therefore, following the same reasoning as in the mean model described above, both sex and father were included in the mapping model as covariates of both the mean and the variance. As with the other tests, the genome permutation procedures described in **Statistical Methods** were used to control FWER.

A genomewide significant mQTL was identified on chromosome 11 (Figure 5, blue line). The peak was at 69.6 cM with FWER-adjusted *p* = 0.011, with the closest marker being D11MIT11 at 75.7 cM with FWER-adjusted *p* = 0.016. Nonparametric bootstrap resampling, using 1,000 resamples (after Visscher *et al.* 1996), established a 90% confidence interval for the QTL from 50 to 75 cM. This region overlaps with the “suggestive” region identified in the traditional analysis.

By the traditional definition of percent variance explained, following from a fitting of the standard linear model, this QTL explains 2.1% of phenotype variance. Though, given the variance heterogeneity inherent in the DGLM that was used to detect this QTL, this quantity is better considered the “average” percent variance explained. The ratio of the QTL variance to the sum of QTL variance, covariate variance, and residual variance ranges from 1% to 6% across the population, based on the heterogeneity of residual variance.

#### Understanding the novel QTL

The mQTL on chromosome 11 was identified by the DGLM_M_ test, but not by the standard linear model or Cao’s mQTL test. The additional power of the DGLM_M_ test over these other tests relates to its accommodation of background variance heterogeneity (BVH).

Specifically, the DGLM reweighted each observation based on its residual variance, according to the sex and F1 father of the mouse. This BVH is visually apparent when the residuals from the standard linear model are plotted, separated out by father (Figure 6).

**Figure 6.**
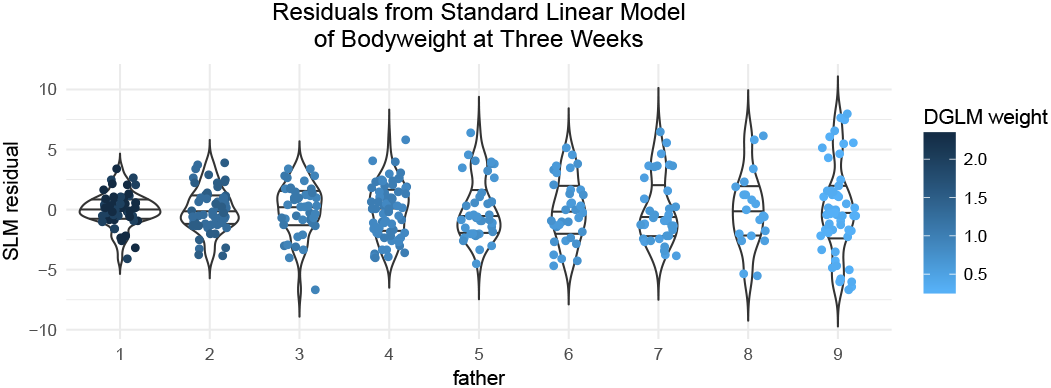
Residuals from the standard linear model for body weight at three weeks, with sex and father as covariates, stratified by father. It is evident that fathers differed in the residual variance of the offspring they produced. For example, the residual variance of offspring from father 1 is greater than that of father 2 and 7. Here, points are colored by their predicted residual variance in the fitted DGLM with sex and father as mean and variance covariates.

Some fathers, for example fathers 2 and 7, appear to have offspring with less residual variance than average, whereas others, for example father 1, seem to have offspring with more residual variance than average. The DGLM captured these patterns of variance heterogeneity, and estimated the effect of each father on the log standard deviation of the observations (Figure 7). Based on these estimated variance effects, observations were upweighted (e.g. fathers 2 and 7) and downweighted (e.g. father 1). This weighting gave the DGLM-based mapping approach more power to reject the null as compared with the SLM.

**Figure 7.**
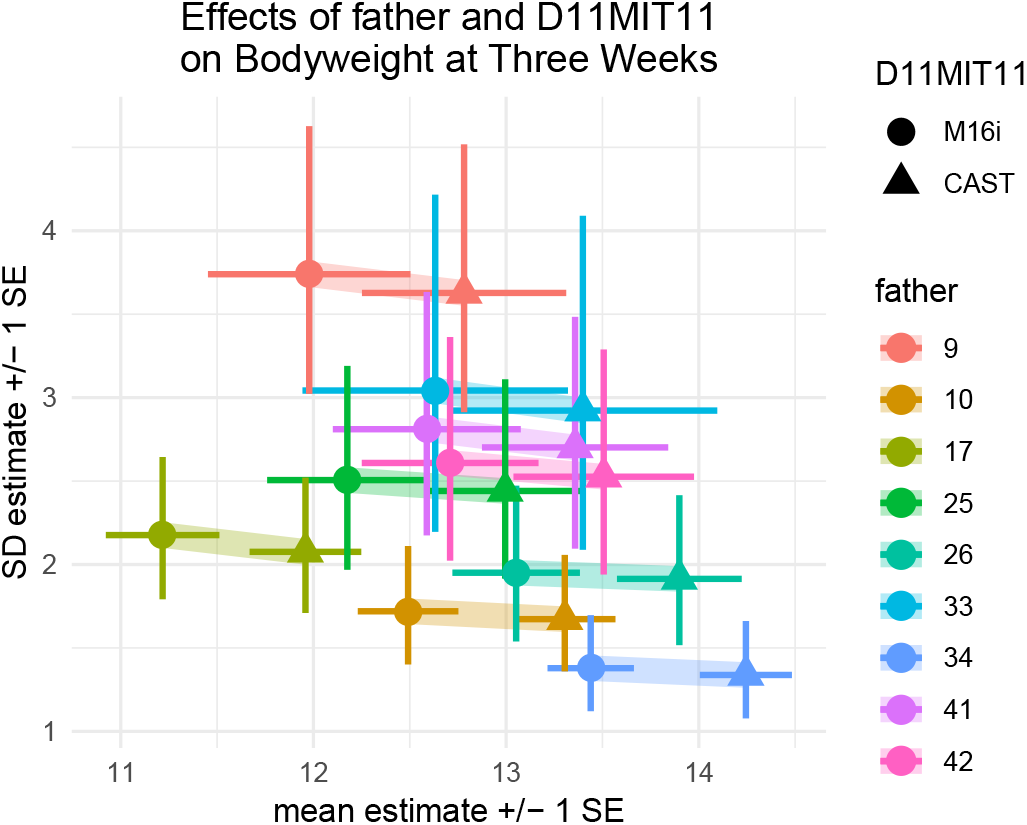
The predictive mean and standard deviation of mice in the mapping population based on father and genotype at the top marker, D11MIT11 on chromosome 11. The genotype effect, illustrated by the colored ribbons is almost entirely horizontal, indicating a difference in means across genotype groups but no difference in variance, consistent with the identification of this QTL as a pure mQTL. The father effects, illustrated by the spread of colored crossbars, have both mean and variance components. For example, father 7 (blue) has the highest predictive mean and lowest predictive standard deviation. His offspring were upweighted in the QTL analysis based on their low standard deviation. Father 1 (red) has an average predictive mean and the highest predictive standard deviation. His offspring were downweighted in the QTL analysis based on their high standard deviation. Note: the effect of sex on phenotype mean and variance was modeled, then marginalized out for readability.

#### Other phenotypes

For brevity, we described in detail only the results of the DGLM-based analysis of body weight at three weeks; but, of the eight phenotypes from this cross available on the Mouse Phenome Database, the mean-variance approach to QTL mapping discovered new QTL in four. Five of the eight phenotypes — body weight at twelve days, three weeks, and six weeks, as well as subcutaneous and gonadal fat pad thickness — exhibited BVH due to father, and for each we performed both traditional QTL mapping using the SLM and mean-variance QTL mapping using the DGLM. This reweighting changed the results in three cases: For body weight at three weeks (Figure S15) and six weeks (Figure S16), we identified one new mQTL and two new vQTL respectively. For subcutaneous fat pad thickness, we discovered one mQTL and “undiscovered” one mQTL (Figure S17). That is, after reweighting the observations based on the observed variance of each father, one locus that was overlooked by SLM was identified as an mQTL and one locus that was identified by SLM as an mQTL was no longer found to have a statistically significant association with the phenotype.

## DISCUSSION

Since the recognition that variance effects can be attributable to individual genes, a growing body of research has asked questions about the prevalence of such effects (Huang *et al.* 2015), their evolutionary origins (canalization, robustness), their ramifications (decanalization in disease, increased variation) (Gibson 2009; Fre*und et al.* 2013; Lin *et al.* 2016), and how the identification of such genes can provide a signal of, and thereby serve as a route to identify, higher order interactions such as epistasis or GxE (Struchalin *et al.* 2010; Rönnegård and Valdar 2012; Forsberg and Carlborg 2017). These studies have promoted detection of variance heterogeneity as path to new biological discovery. But less attention has been paid to this corollary: if a phenotype is subject to variance-controlling factors, then, whether or not identifying those factors is of direct interest, they will induce background variance heterogeneity that can affect inference of more standard targets, including mean-affecting QTL. In other words, interest in identifying sources of BVH may be of most interest to a subset of researchers, but interest in accommodating it should be more widespread.

Our simulation studies showed that modeling BVH when it is present increases power to detect mQTL, vQTL and mvQTL. Our reanalysis of the Leamy et al. dataset demonstrated that accommodating BVH can lead to detection of mQTL that would otherwise be overlooked.

In both cases, of the methods compared, the most powerful were those based on the DGLM, with the most robust version of that using the locusperm significance procedure. These results should not be too surprising. The DGLM was the only method examined that can accommodates variance effects arising from both the locus and from other covariates; and the locusperm method (and genomeperm, its genomewide analog) is least reliant on parametric assumptions. We would expect other methods that allow flexible modeling of covariate effects on variance to be competitive in these regards, *e.g.*, the recent Bayesian hierarchical model of Dumitrascu *et al.* (2018).

Beyond advocating any particular method, however, our results can be used to draw attention to a number of more general points about 1) the relationship between increased residual variance, observation weighting and downstream inference; 2) how knowledge of variance effects can be exploited in experimental design, analysis and reanalysis; 3) the sensitivity of variance effect detection to distributional assumptions and how strategies such as permutation or variable transformation can mitigate this; and, 4) the challenges in reporting quantitative genetic parameters under heteroskedasticity.

### Residual variance, weighting, and inference for mean effect QTL

The additional power of mean-variance QTL mapping to detect mQTL in general—and of DGLM_M_ to detect mQTL in the presence of BVH in particular—can be seen as deriving from how data is reweighted. Consider heteroskedastic data modeled as *y_i_* ~ N(*mi*, *σ*^2^/*w_i_*), with known weights *w*_1_, …, *w_n_* and known baseline variance *σ*^2^. The log-likelihood can be written as *ℓ* = const WRSS/2*σ*^2^, such that the key quantity to be minimized in a maximum likelihood fit is

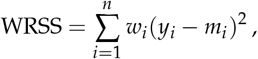

the weighted residual sum of squares, that is, the squared discrepancies between the observed phenotype *yi* and its predicted value *mi*, weighted by *wi*. The weights therefore affect how much, relatively speaking, each data point contributes to the likelihood: highly imprecise measurements, such as from individuals whose phenotypes are expected to have high variance, have low weight and diminished contribution, whereas as more precise measurements are correspondingly upweighted. In the DGLM, the weight of each observation is determined in the model-fitting process based on the phenotype, the experimental covariates, and the QTL genotype, as *w_i_* = *e^−υi^*. In the SLM, weights can be specified, but they cannot be co-estimated with covariate and QTL effects. The improvement of the DGLM over the SLM and CaoM under BVH stems entirely from its greater ability to capture this additional information, and thereby give more credence to phenotype values that are more precise.

We note a related approach to correcting inference of mean effects in the face of heteroskedasticity not considered here is the use of heteroskedastic consistent covariance matrix estimators (HCCMEs) [Long and Ervin (2000) and refs therein]. Also known as “sandwich” estimators, these use estimated residuals from the SLM to characterize heteroskedasticity empirically and thereby estimate adjusted, heteroskedastic-consistent versions of the effect standard errors. Importantly, HCCMEs do not require the source of heteroskedasticity to be identified, and they have seen recent use in genetic association [*e.g.*, Barton *et al.* (2013); Rao and Province (2016)]. However, this comes at a cost: when a variable that does predict heteroskedasticity can be identified, HCCMEs will tend to be inefficient compared with a model-based estimator (Wakefield 2013), such as the DGLM.

### Implications for experimental design, analysis and reanalysis

The possibility that some individuals could be predictably more variable than others has clear implications for experimental design. A key parameter in the design of experiments is the number of replicates, typically specified to provide adequate precision of, and thereby power to detect, an estimated effect. But foreknowledge that residual variance will differ for certain groups suggests a more nuanced approach that explicitly weighs replicates against intrinsic variability.

For example, when designing an experiment on a population that happens to have a known, segregating vQTL that is not itself the focus of interest but would induce BVH, it may be preferable to allocate a disproportionate share of the replication to individuals in the high-variability genotype class. In such cases, it then becomes additionally helpful to understand at what level(s) the heterogeneous variance manifests. Specifically, increased variability could arise from greater between-individual variation or greater within-individual variation [cf more levels of variability described in Table 1 of Rönnegård and Valdar (2011)]; whereas the between-individual case warrants additional biological replicates, the within-individual case could be addressable (potentially more cheaply) with additional technical replicates.

Alternatively, the recognition that some individuals are predictably high variance may be a reason to exclude them entirely, or, more generally, to opt for conditions and population subsets for which residual variance is predicted to be minimal. If such a variance-minimizing population can be achieved without changing the genetic effects present, it would have an improved signal-to-noise ratio and provide better power to detect genetic effects.

A more standard situation is that a vQTL (or other BVH factor) is not recognized until the experiment is first analyzed. In this case, it would make sense to perform a re-analysis, with the vQTL included as a variance-affecting covariate. Doing so should increase power to detect both mQTL and other vQTL.

### vQTL mapping: pros and cons of the rank inverse normal transformation

The presence of BVH can be disruptive to a test for a vQTL. A simplistic test compares a heteroskedastic alternative model with a homoskedastic null. BVH confuses the comparison by making the true null heteroskedastic. In doing so, it increases the false positive rate for asymptotic tests that disregard BVH and reduces power when FPR is empirically controlled (see, *e.g.*, CaoV results in Table 3).

In this context it is therefore interesting to consider the crude— but often used—device of the rank inverse normal transformation. The RINT reshapes away any kurtosis (fatter tails), a key signature of heteroskedasticity, without any reference to its source. As such, it is logical that in the detection of vQTL it would have both beneficial and harmful properties.

In the case where there is no known driver of BVH, represented by the simulations examining CaoV, the RINT procedure acts as an insurance policy: if there truly is no BVH, the test suffers a modest decrease in power; but if there truly is BVH from an unknown source, it averts the drastic FPR inflation under the standard (*i.e.*, non-empirical) p-value procedure.

In the case where researchers are confident that, after correcting for known BVH drivers, the residuals are homoskedastic (represented by the DGLMV simulations), the RINT procedure is unnecessary, costing power with its conservatism in the absence of BVH and paradoxically creating even more conservative behavior in the presence of BVH.

The aforementioned disadvantages of RINT assume the phenotype data has an underlying normal distribution, either as given or after a deducible transformation [*e.g.*, via the Box-Cox procedure or similar; (Box and Cox 1964)]. When the data is highly non-normal, however, both the RINT and the locusperm procedure would provide valid inference, and perhaps the most robust approach would be to use the two in combination. Nonetheless, where normality approximately holds, whether as given or after a simple transformation, we strongly prefer the locusperm procedure without RINT: across all simulation scenarios it exhibited at worst slight conservatism when applied to DGLM-based tests and represents a useful step toward FWER control.

### Permutation schemes for other populations

Our preferred permutation scheme, locusperm (or its genomewide equivalent, genomeperm), is applicable to populations in which genotypes under the null are exchangeable. As such, it holds not only for F2 and backcrosses but also, for example, in equally-related recombinant inbred line panels such as the Collaborative Cross and another similar replicable multiparent populations. For example, in the (mQTL) study of Mosedale *et al.* (2017), the use of locus genotypes (or genotype probabilities) would simply be replaced by founder haplotypes that could then be randomly exchanged across lines.

In non-exchangeable populations, however, such as those requiring polygenic random effect terms [*e.g.*, Kennedy *et al.* (1992)], although the DGLM could be applied via its random effects generalization, DHGLM (Felleki *et al.* 2012), the permutation scheme may need revision. In particular, a permutation scheme in which all permutations are equally likely may not comport with a reasonable null, and it may be more appropriate to allocate higher probabilities to permutations that preserve overall genetic similarity (Abney 2015; Roach and Valdar 2018; Berrett *et al.* 2018). Although we not have a specific solution, we suspect that the necessity of such revisions, at least for the DGLMV test, will depend on the extent to which the observed heteroskedasticity is polygenic.

### Percent variance explained

Variance heterogeneity complicates the notion of percent variance explained (PVE) by a QTL. Assuming the QTL has the same effect on the expected value of the phenotype of all individuals, it will explain a larger percent of total variance for individuals with lower than average residual variance, and vice versa for individuals with higher than average residual variance. In light of this observation, the percent variance explained can either be reported as “average percent variance explained” or can be calculated for some representative sub-groups. For example, if there is variance heterogeneity across sexes, it would be reasonable to report the PVE of a QTL for both males and females, or if a vQTL is known to be present elsewhere in the genome, report the PVE for each vQTL genotype as in Yang *et al.* (2012).

### Guidelines for detecting and QTL mapping in the presence of BVH

To select the right test and procedure to assess significance, it is important to establish whether there is any BVH present. We advocate fitting the DGLM with all potential BVH drivers as variance covariates, then including any that are statistically significant as variance covariates in the mapping model to improve power to detect QTL. Then, given that

1. The DGLM-based tests dominate all other tests in the presence of BVH,
2. the locusperm procedure accurately controls the FPR of the DGLM-based tests in the presence of BVH, whether the source is known or not, and
3. the locusperm procedure can be extended into the genomeperm procedure to control FWER, we advocate for the analysis of experimental crosses that exhibit BVH with the three DGLM-based tests (DGLMM, DGLMV, and DGLMMV) and, where the individuals in the population are exchangeable (as in an F2 or backcross) or where partial exchange-ability can be suitably identified [*e.g.*, see (Churchill and Doerge 1994; Zou *et al.* 2006; Churchill and Doerge 2008)], the use of our described genomeperm procedures, which permute the genome in selective parts of the model, to assess genomewide significance.

Because this procedure involves three families of tests rather than one family as would be typical with an SLM-based analysis, an additional correction may be desired to control experiment-wise error rate. DGLMM and DGLMV are orthogonal tests (Smyth 1989), but DGLMMV is neither orthogonal nor identical to either, so the effective number of families is between two and three. One reasonable, heuristic approach to control experiment-wise error rate is simply to lower the acceptable FWER, *e.g.* replacing the standard 0.05 with 0.02.

### Conclusion

In summary, we demonstrate the effect of BVH on QTL mapping of both mQTL and vQTL, and the value of accommodating it using the DGLM. In doing so, we propose a standard protocol for mapping mQTL, vQTL and mvQTL in standard genetics crosses.

## ACKNOWLEDGMENTS

This work was funded by National Institutes of General Medical Sciences grants R01-GM104125 (RWC,WV), R35-GM127000 (RWC,WV), T32-GM067553 (RWC); National Heart, Lung and Blood Institute grant R21 HL126045 (RWC,WV); National Library of Medicine grant T32-LM012420 (RWC); and a National Institute of Mental Health grant F30-MH108265 (RWC). The authors thank Larry Leamy for helpful comments.

## APPENDIX Calculation of an additive effect to explain a given proportion of total variance in an F2 intercross

The variance attributable to a genetic factor with alleles (AA, AB, BB) at frequency (0.25, 0.5, 0.25), additive effect *a* and no dominance effect is:

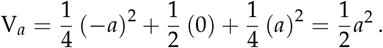

For a genetic factor that explains a fraction *p* of total phenotype variance:

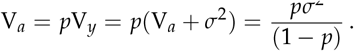

Combining and solving for *a* gives 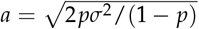.

## SUPPLEMENTARY MATERIALS

### Simulation Details

In simulation with BVH present, the group-wise effects on the log standard deviation were ***γ*** = [0.4, 0.2, 0, 0.2, 0.4]. Though ***γ̄*** = 0, the exponential transform connecting these effects to the standard deviation results in a simulated phenotype with slightly more total variance than one without BVH. Therefore, the additive effect of the locus on phenotype mean was adjusted when BVH was introduced, in order to maintain a constant percent variance explained by the mean effect. The following values were used in the simulation.

|  | no BVH | yes BVH |
| --- | --- | --- |
| null | $\alpha = 0, \theta = 0$ | $\alpha = 0, \theta = 0$ |
| mQTL | $\alpha = 0.22, \theta = 0$ | $\alpha = 0.25, \theta = 0$ |
| vQTL | $\alpha = 0, \theta = 0.17$ | $\alpha = 0, \theta = 0.17$ |
| mvQTL | $\alpha = 0.18, \theta = 0.14$ | $\alpha = 0.2, \theta = 0.136$ |

***null locus and mQTL in the absence of BVH:*** All observations have standard deviation 1.
***vQTL in the absence of BVH:*** The genotype-wise standard deviations implied by the additive effect of 0.17 on the log standard deviation are approximately: [0.84, 1.00, 1.19].
***mvQTL in the absence of BVH:*** The genotype-wise standard deviations implied by the additive effect of 0.14 on the log standard deviation are approximately: [0.87, 1.00, 1.15].
***null locus and mQTL in the presence of BVH:*** The covariate-wise standard deviations implied by the effects of [−0.4, −0.2, 0, 0.2, 0.4] on the log standard deviation are approximately: [0.67, 0.82, 1.00, 1.22, 1.49].
***vQTL in the presence of BVH:*** Locus and covariate effects on the residual variance combine additively on the log standard deviation scale, yielding 15 distinct standard deviations:

***mvQTL in the presence of BVH:*** Locus and covariate effects on the residual variance combine additively on the log standard deviation scale, yielding 15 distinct standard deviations:

|  | genotype |  |  |
| --- | --- | --- | --- |
| covar | -1 | 0 | 1 |
| 1 | 0.57 | 0.67 | 0.79 |
| 2 | 0.69 | 0.82 | 0.97 |
| 3 | 0.84 | 1.00 | 1.19 |
| 4 | 1.03 | 1.22 | 1.45 |
| 5 | 1.26 | 1.49 | 1.77 |

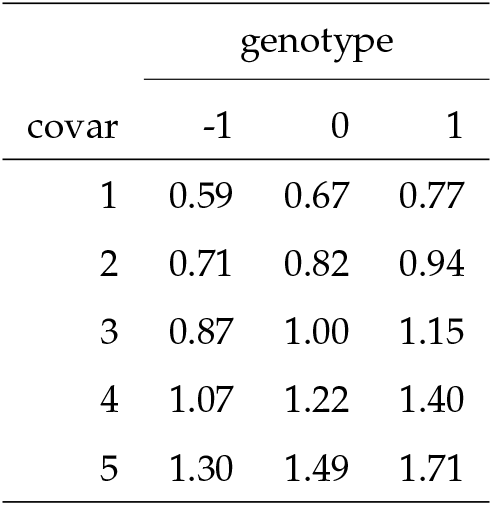

### AUC Table

**Table S1.** Area under the curve for the six non-null scenarios and all 32 test-procedures.

| test | procedure | BVH absent |  |  | BVH present |  |  |
| --- | --- | --- | --- | --- | --- | --- | --- |
|  |  | mQTL | vQTL | mvQTL | mQTL | vQTL | mvQTL |
| SLM | standard | 0.931 | 0.502 | 0.860 | 0.926 | 0.495 | 0.858 |
|  | RINT | 0.930 | 0.499 | 0.855 | 0.935 | 0.493 | 0.866 |
|  | residperm | 0.930 | 0.503 | 0.857 | 0.928 | 0.495 | 0.857 |
|  | locusperm | 0.931 | 0.503 | 0.858 | 0.927 | 0.494 | 0.858 |
| Cao<sub>M</sub> | standard | 0.930 | 0.504 | 0.862 | 0.930 | 0.499 | 0.861 |
|  | RINT | 0.929 | 0.497 | 0.858 | 0.933 | 0.495 | 0.865 |
|  | residperm | 0.930 | 0.502 | 0.861 | 0.926 | 0.494 | 0.860 |
|  | locusperm | 0.929 | 0.501 | 0.859 | 0.926 | 0.495 | 0.859 |
| DGLM<sub>M</sub> | standard | 0.929 | 0.514 | 0.863 | 0.964 | 0.503 | 0.914 |
|  | RINT | 0.929 | 0.510 | 0.858 | 0.963 | 0.506 | 0.911 |
|  | residperm | 0.925 | 0.503 | 0.856 | 0.962 | 0.496 | 0.908 |
|  | locusperm | 0.925 | 0.500 | 0.854 | 0.961 | 0.493 | 0.906 |
| Levene's test | standard | 0.500 | 0.912 | 0.836 | 0.489 | 0.887 | 0.804 |
|  | RINT | 0.486 | 0.908 | 0.818 | 0.485 | 0.877 | 0.782 |
|  | residperm | 0.502 | 0.916 | 0.842 | 0.497 | 0.890 | 0.809 |
|  | locusperm | 0.502 | 0.916 | 0.841 | 0.496 | 0.890 | 0.808 |
| Cao<sub>V</sub> | standard | 0.501 | 0.937 | 0.870 | 0.593 | 0.930 | 0.864 |
|  | RINT | 0.485 | 0.925 | 0.844 | 0.495 | 0.882 | 0.795 |
|  | residperm | 0.500 | 0.933 | 0.864 | 0.503 | 0.887 | 0.802 |
|  | locusperm | 0.497 | 0.932 | 0.863 | 0.497 | 0.883 | 0.802 |
| DGLM<sub>V</sub> | standard | 0.503 | 0.933 | 0.864 | 0.503 | 0.937 | 0.864 |
|  | RINT | 0.491 | 0.919 | 0.838 | 0.440 | 0.891 | 0.799 |
|  | residperm | 0.501 | 0.928 | 0.859 | 0.417 | 0.887 | 0.797 |
|  | locusperm | 0.501 | 0.928 | 0.856 | 0.497 | 0.931 | 0.858 |
| Cao<sub>MV</sub> | standard | 0.900 | 0.908 | 0.941 | 0.910 | 0.901 | 0.940 |
|  | RINT | 0.897 | 0.890 | 0.930 | 0.896 | 0.838 | 0.918 |
|  | residperm | 0.896 | 0.902 | 0.938 | 0.871 | 0.858 | 0.911 |
|  | locusperm | 0.897 | 0.904 | 0.938 | 0.873 | 0.859 | 0.911 |
| DGLM<sub>MV</sub> | standard | 0.900 | 0.901 | 0.938 | 0.941 | 0.905 | 0.959 |
|  | RINT | 0.893 | 0.887 | 0.929 | 0.933 | 0.849 | 0.941 |
|  | residperm | 0.892 | 0.893 | 0.933 | 0.912 | 0.856 | 0.933 |
|  | locusperm | 0.894 | 0.894 | 0.933 | 0.938 | 0.897 | 0.955 |

### Standard Error Table

**Table S2.** Standard errors for the values in Table S2.

| test | procedure | BVH absent |  |  |  | BVH present |  |  |  |
| --- | --- | --- | --- | --- | --- | --- | --- | --- | --- |
|  |  | null | mQTL | vQTL | mvQTL | null | mQTL | vQTL | mvQTL |
| SLM | standard | 0.0044 | 0.0089 | 0.0045 | 0.0098 | 0.0044 | 0.0090 | 0.0044 | 0.0098 |
|  | RINT | 0.0043 | 0.0089 | 0.0044 | 0.0098 | 0.0044 | 0.0089 | 0.0043 | 0.0098 |
|  | residperm | 0.0043 | 0.0089 | 0.0044 | 0.0098 | 0.0043 | 0.0090 | 0.0044 | 0.0098 |
|  | locusperm | 0.0043 | 0.0090 | 0.0044 | 0.0098 | 0.0044 | 0.0090 | 0.0044 | 0.0098 |
| Cao<sub>M</sub> | standard | 0.0044 | 0.0089 | 0.0044 | 0.0098 | 0.0045 | 0.0090 | 0.0043 | 0.0098 |
|  | RINT | 0.0044 | 0.0089 | 0.0042 | 0.0098 | 0.0045 | 0.0089 | 0.0043 | 0.0098 |
|  | residperm | 0.0044 | 0.0090 | 0.0043 | 0.0098 | 0.0044 | 0.0091 | 0.0042 | 0.0098 |
|  | locusperm | 0.0043 | 0.0090 | 0.0042 | 0.0098 | 0.0043 | 0.0091 | 0.0042 | 0.0098 |
| DGLM<sub>M</sub> | standard | 0.0046 | 0.0089 | 0.0045 | 0.0098 | 0.0047 | 0.0073 | 0.0046 | 0.0094 |
|  | RINT | 0.0045 | 0.0089 | 0.0045 | 0.0098 | 0.0046 | 0.0073 | 0.0045 | 0.0094 |
|  | residperm | 0.0043 | 0.0091 | 0.0042 | 0.0098 | 0.0044 | 0.0075 | 0.0043 | 0.0095 |
|  | locusperm | 0.0043 | 0.0091 | 0.0042 | 0.0098 | 0.0043 | 0.0076 | 0.0043 | 0.0095 |
| Levene's test | standard | 0.0041 | 0.0042 | 0.0094 | 0.0098 | 0.0043 | 0.0041 | 0.0098 | 0.0096 |
|  | RINT | 0.0041 | 0.0039 | 0.0095 | 0.0097 | 0.0042 | 0.0039 | 0.0098 | 0.0093 |
|  | residperm | 0.0043 | 0.0044 | 0.0093 | 0.0098 | 0.0044 | 0.0043 | 0.0097 | 0.0096 |
|  | locusperm | 0.0042 | 0.0044 | 0.0093 | 0.0098 | 0.0044 | 0.0043 | 0.0097 | 0.0096 |
| Cao<sub>V</sub> | standard | 0.0045 | 0.0044 | 0.0087 | 0.0098 | 0.0067 | 0.0067 | 0.0087 | 0.0098 |
|  | RINT | 0.0041 | 0.0040 | 0.0091 | 0.0098 | 0.0042 | 0.0042 | 0.0098 | 0.0095 |
|  | residperm | 0.0043 | 0.0043 | 0.0088 | 0.0098 | 0.0044 | 0.0043 | 0.0098 | 0.0096 |
|  | locusperm | 0.0043 | 0.0042 | 0.0089 | 0.0098 | 0.0044 | 0.0042 | 0.0098 | 0.0096 |
| DGLM<sub>V</sub> | standard | 0.0045 | 0.0044 | 0.0088 | 0.0098 | 0.0046 | 0.0043 | 0.0087 | 0.0098 |
|  | RINT | 0.0041 | 0.0039 | 0.0092 | 0.0098 | 0.0030 | 0.0029 | 0.0098 | 0.0094 |
|  | residperm | 0.0043 | 0.0043 | 0.0090 | 0.0098 | 0.0028 | 0.0027 | 0.0098 | 0.0093 |
|  | locusperm | 0.0043 | 0.0043 | 0.0090 | 0.0098 | 0.0044 | 0.0042 | 0.0090 | 0.0098 |
| Cao<sub>MV</sub> | standard | 0.0044 | 0.0096 | 0.0095 | 0.0086 | 0.0062 | 0.0094 | 0.0094 | 0.0085 |
|  | RINT | 0.0042 | 0.0097 | 0.0098 | 0.0090 | 0.0043 | 0.0096 | 0.0098 | 0.0094 |
|  | residperm | 0.0043 | 0.0097 | 0.0096 | 0.0088 | 0.0045 | 0.0098 | 0.0098 | 0.0095 |
|  | locusperm | 0.0043 | 0.0097 | 0.0096 | 0.0088 | 0.0044 | 0.0098 | 0.0098 | 0.0095 |
| DGLM<sub>MV</sub> | standard | 0.0046 | 0.0096 | 0.0096 | 0.0087 | 0.0047 | 0.0086 | 0.0095 | 0.0077 |
|  | RINT | 0.0043 | 0.0097 | 0.0098 | 0.0091 | 0.0038 | 0.0088 | 0.0098 | 0.0087 |
|  | residperm | 0.0043 | 0.0097 | 0.0097 | 0.0089 | 0.0031 | 0.0095 | 0.0098 | 0.0091 |
|  | locusperm | 0.0043 | 0.0097 | 0.0097 | 0.0089 | 0.0045 | 0.0087 | 0.0097 | 0.0080 |

### ROC Curves

The receiver operating characteristics (ROC) curve plots the empirical power as a function of the empirical FPR, reflects the ability of a test-procedure combination to discriminate between QTL and null loci. Specifically, for a given test-procedure and cutoff *c*, the FPR is defined as the fraction null simulations in which the nominal p-value *p* was less than *c*. This quantity is best thought of as the “empirical FPR” and is generally superior to the nominal FPR, since the nominal FPR depends on the assumption that the test is appropriately calibrated while the empirical does not. The power is the fraction of non-null experiments in which the nominal *p*-value is less than *c*.

The ROC curve cannot immediately distinguish between tests that accurately control FPR and those that do not. We added a symbol to each ROC curve at the point where *c* = 0.05. In cases where the point falls on the vertical line at FPR = 0.05, it reflects accurate FPR control. In cases where the point falls to the left or right of the vertical line it reflects a conservative or anti-conservative test, respectively.

**Figure S1.**
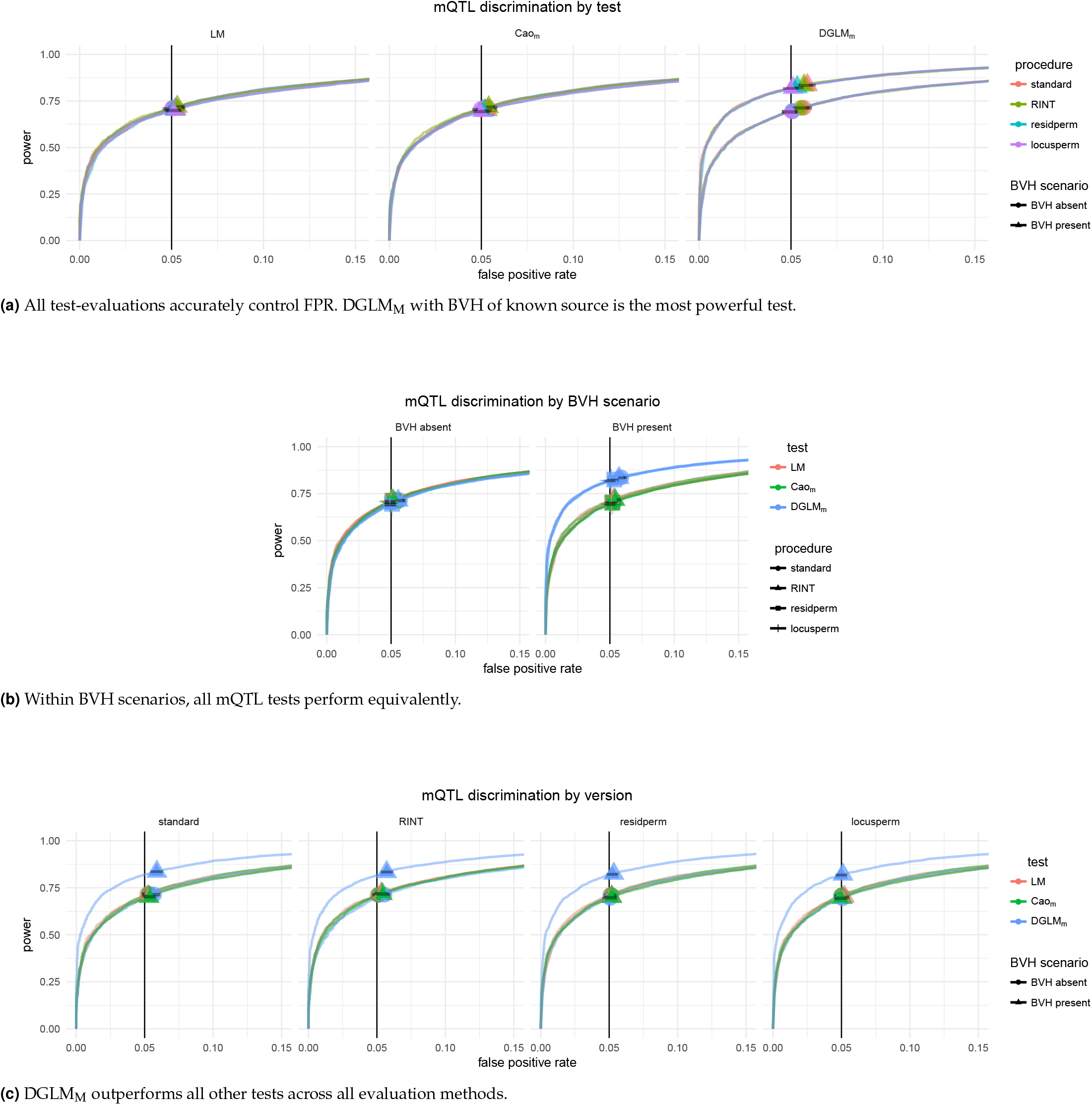
ROC Curves for mQTL tests in the detection of mQTL. The same 28 ROC curves are plotted three times, organized by (a) test, (b) BVH scenario, and (c) significance assessment procedure to allow for comparisons across all dimensions.

**Figure S2.**
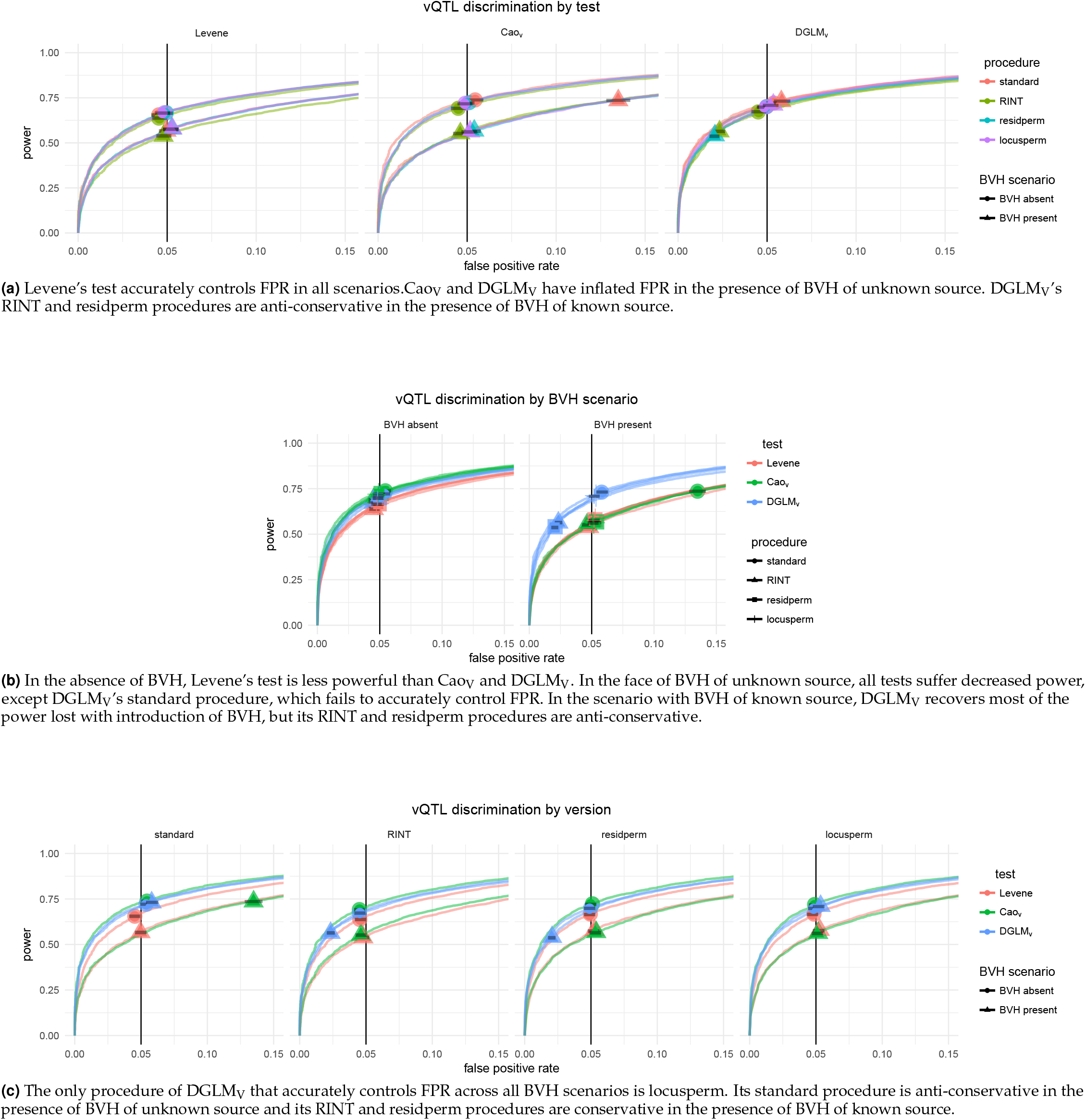
ROC Curves for vQTL tests in the detection of vQTL. The same 28 ROC curves are plotted three times, organized by (a) test, (b) BVH scenario, and (c) significance assessment procedure to allow for comparisons across all dimensions.

**Figure S3.**
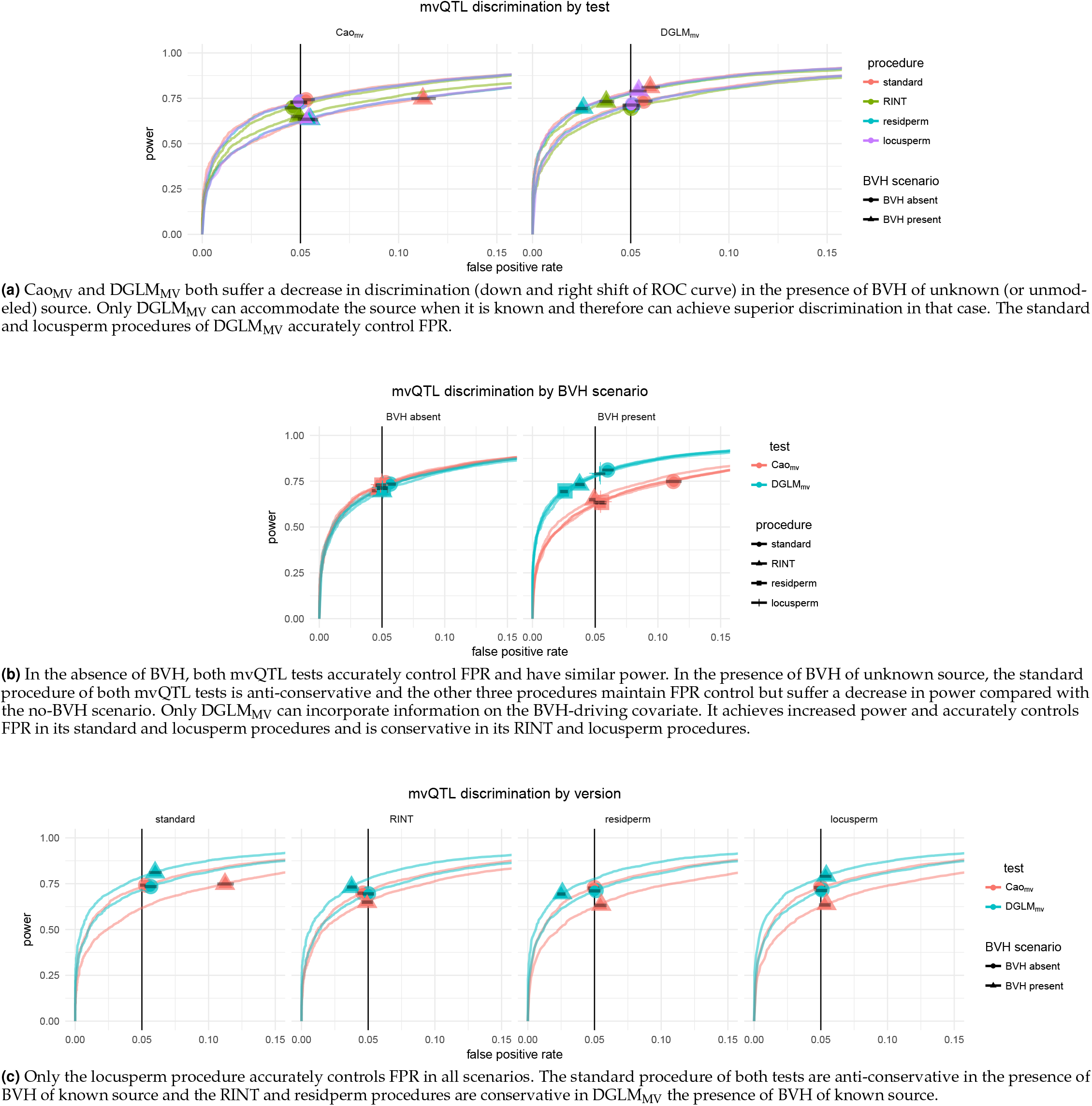
ROC Curves for mvQTL tests in the detection of mvQTL. The same 28 ROC curves are plotted three times, organized by (a) test, (b) BVH scenario, and (c) significance assessment procedure to allow for comparisons across all dimensions.

### QQ Plots

**Figure S4.**
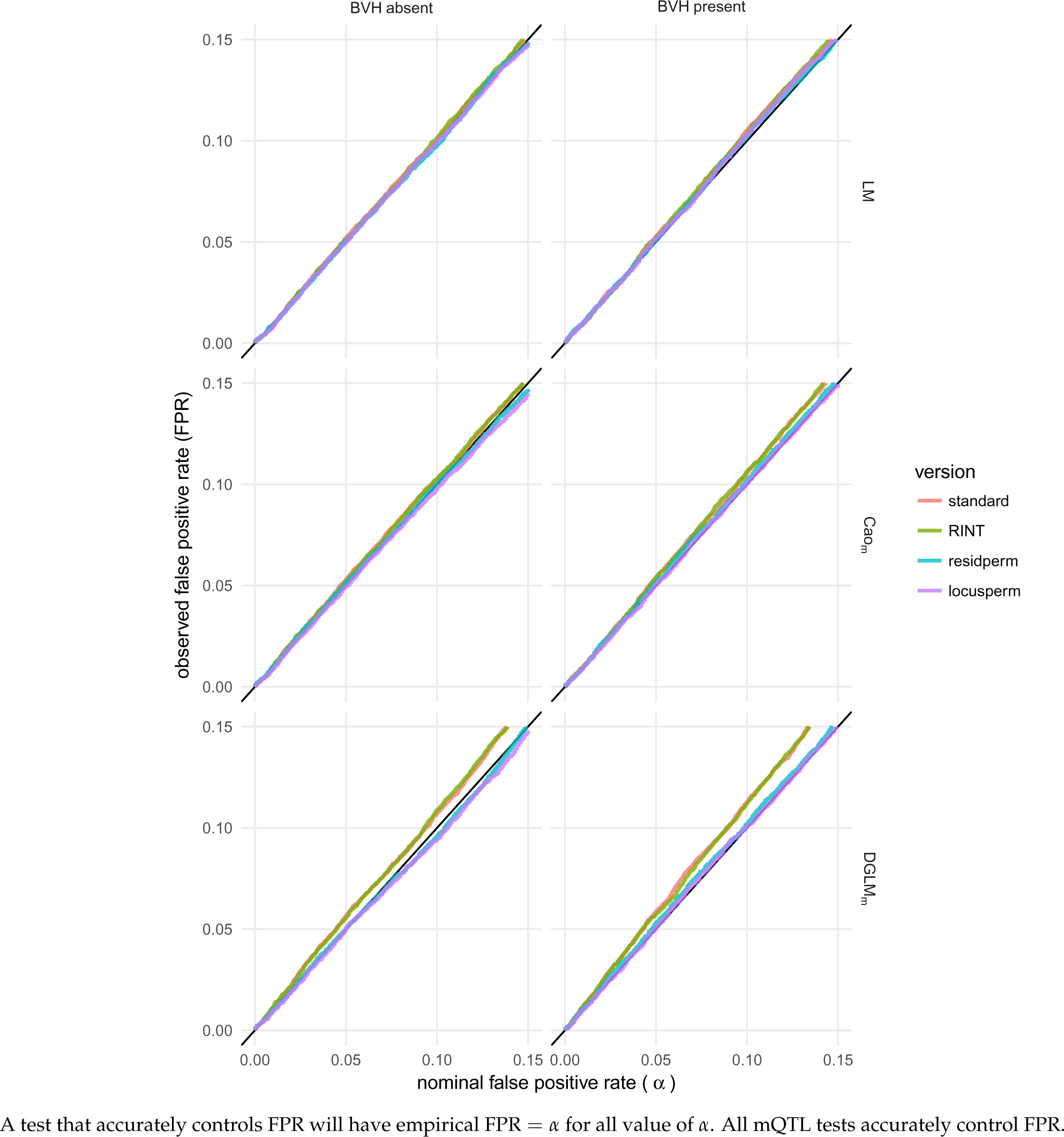
The empirical false positive rate of each test with each procedure for each nominal false positive rate, *α*, in [0, 0.1].

**Figure S5.**
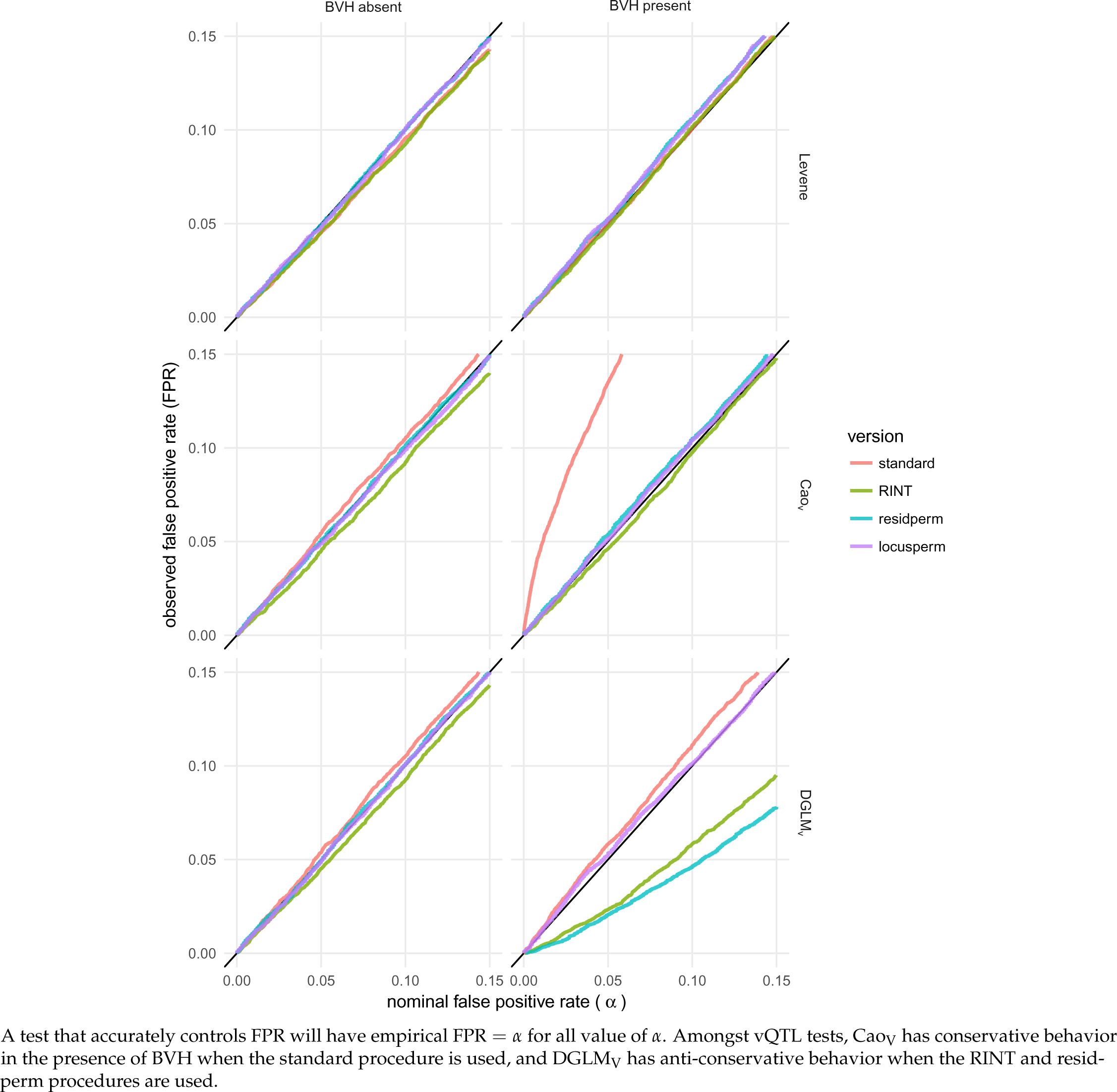
The empirical false positive rate of each test with each procedure for each nominal false positive rate, *α*, in [0, 0.1].

**Figure S6.**
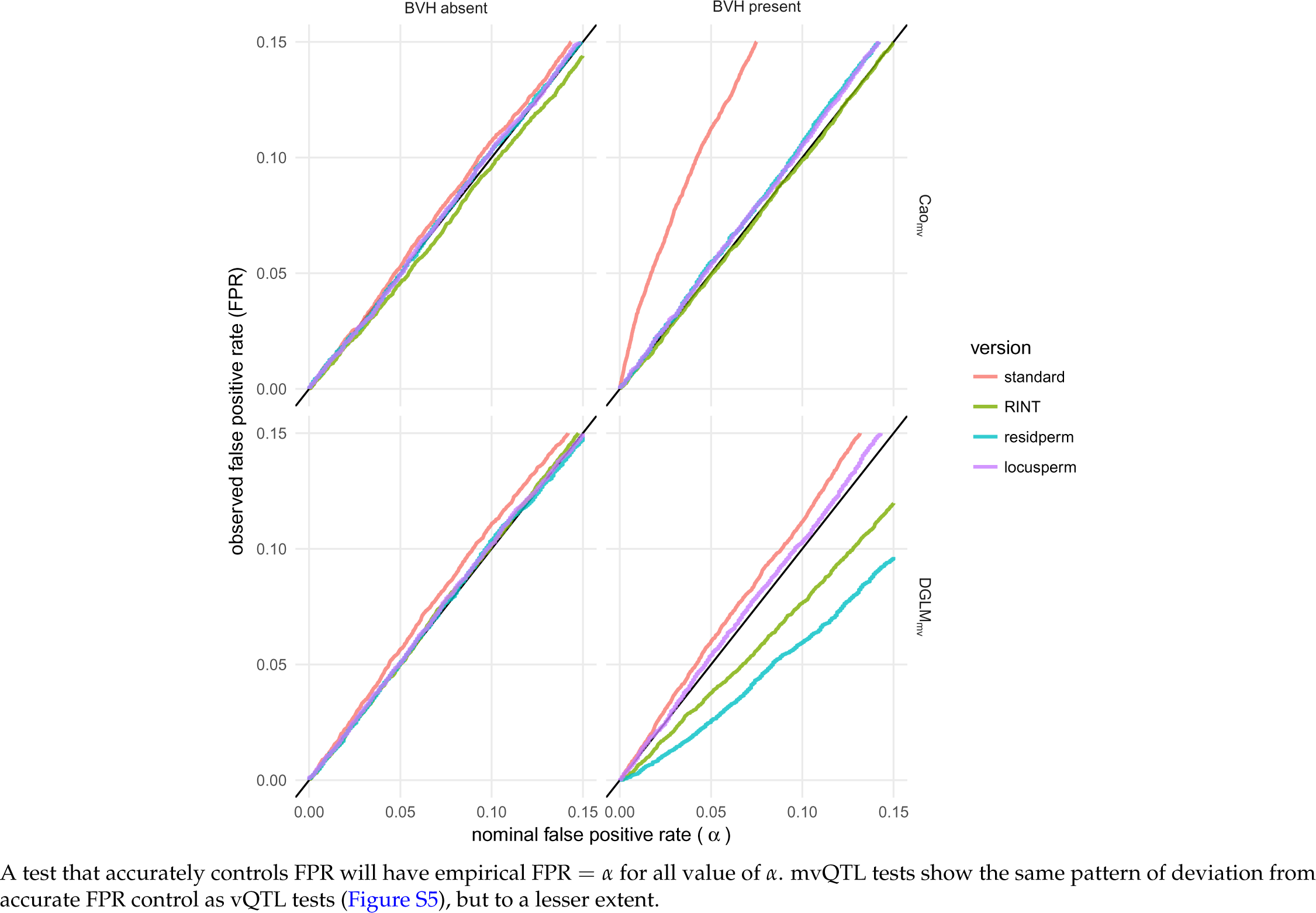
The empirical false positive rate of each test with each procedure for each nominal false positive rate, *α*, in [0, 0.1].

### False positive rate of all test-procedure combinations in all scenarios, caterpillar plot

**Figure S7.**
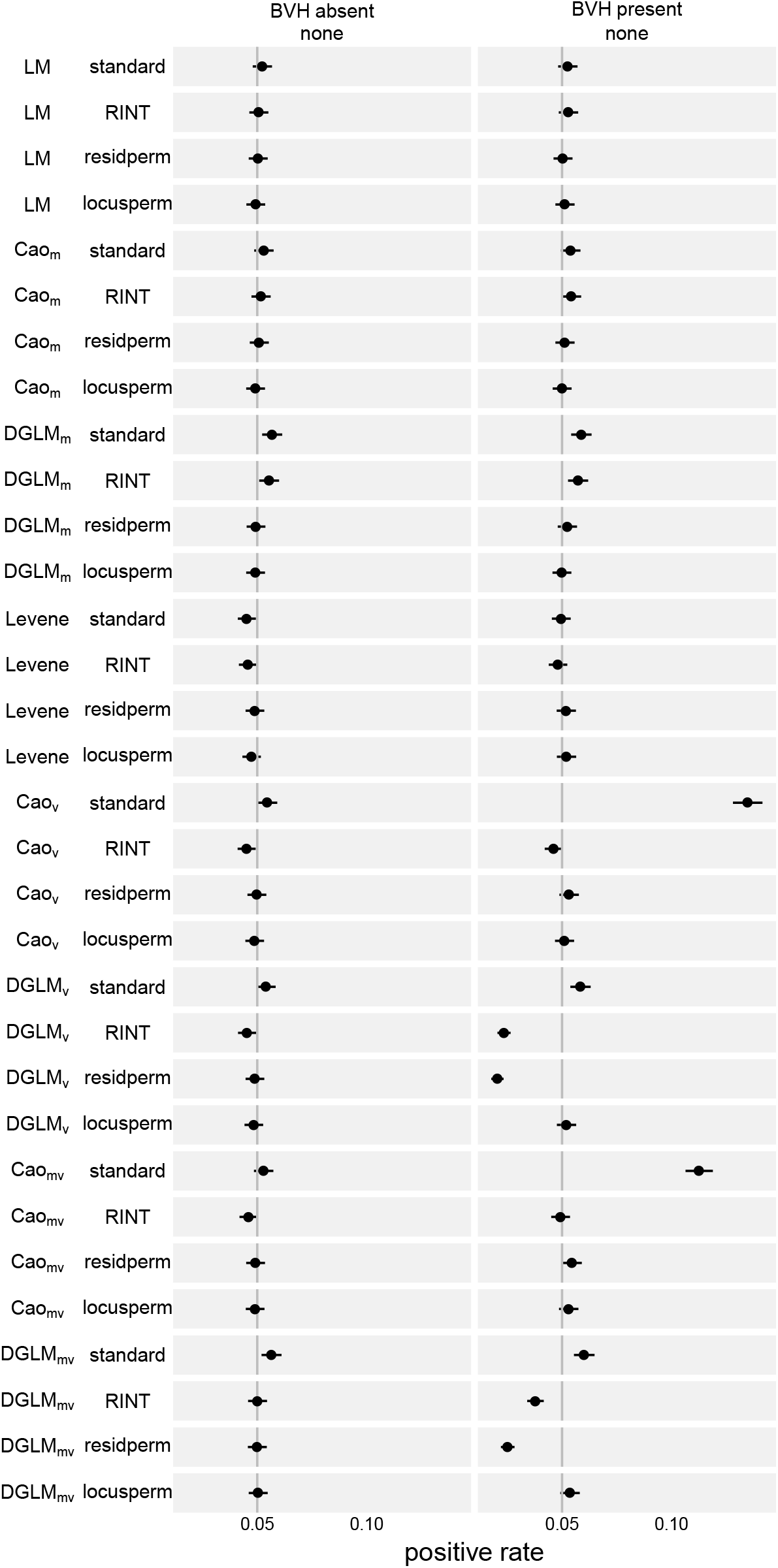

### Cao’s Profile-Likelihood Aprroximation is Extremely Accurate

**Figure S8.**
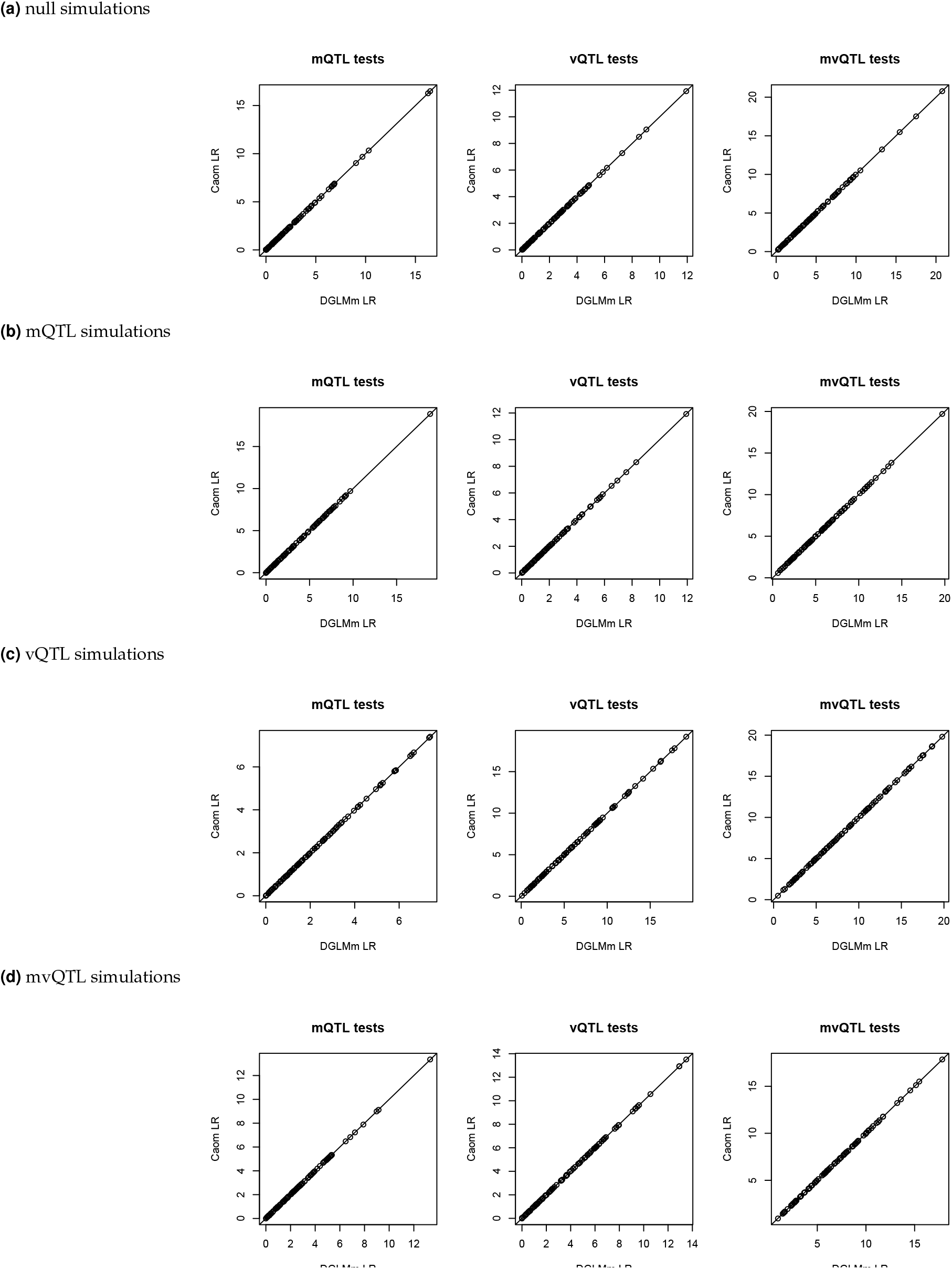
On simulated null loci, mQTL, vQTL, and mvQTL, Cao’s method had identical likelihood ratio to the DGLM without any variance covariates. This result illustrates that although Cao’s method uses a two-step profile likelihood method which is not guaranteed to attain the global maximum likelihood parameter values and the DGLM uses an iterative procedure that does guarantee this attainment, the difference between the two is negligible in this application.

### Cao’s Tests for All Phenotypes with BVH

**Figure S9.**
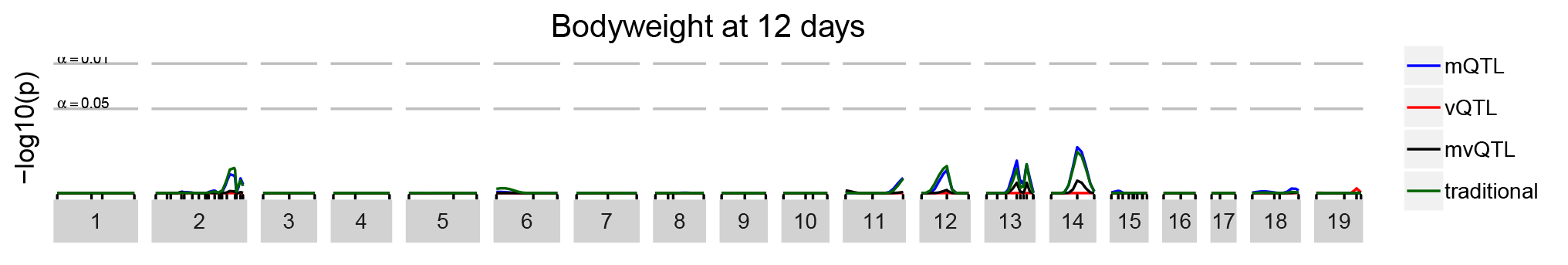
These scans were conducted with the DGLM, without accounting for effects of sex and father on variance, shown by simulation to be identical to Cao’s tests (Figure S8).

**Figure S10.**
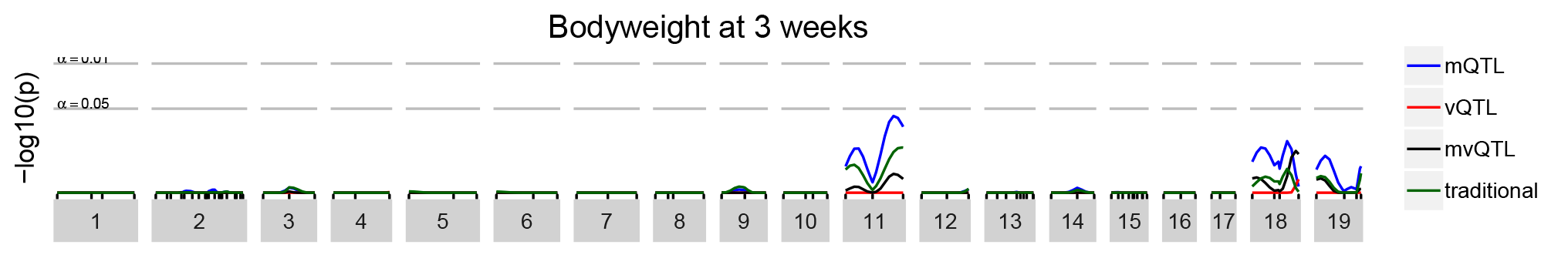

**Figure S11.**
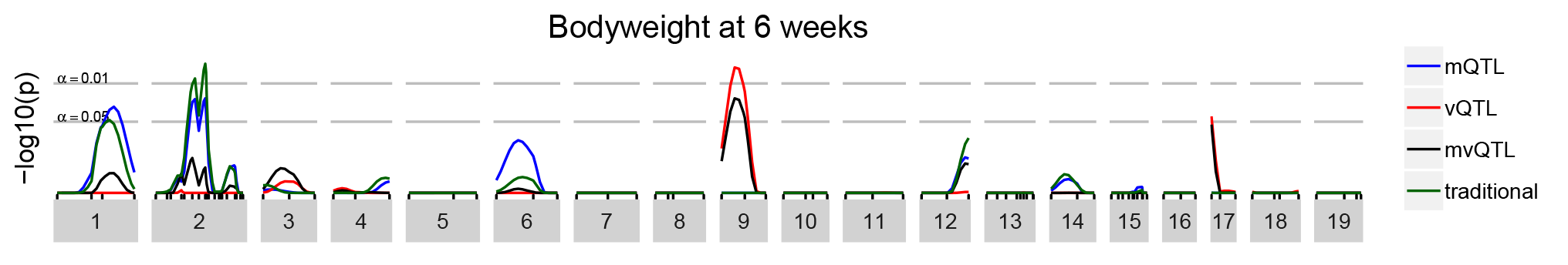

**Figure S12.**
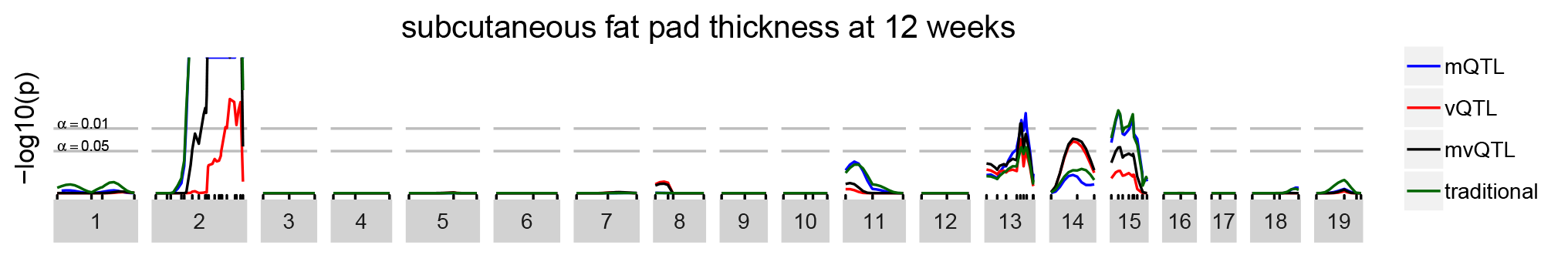

**Figure S13.**
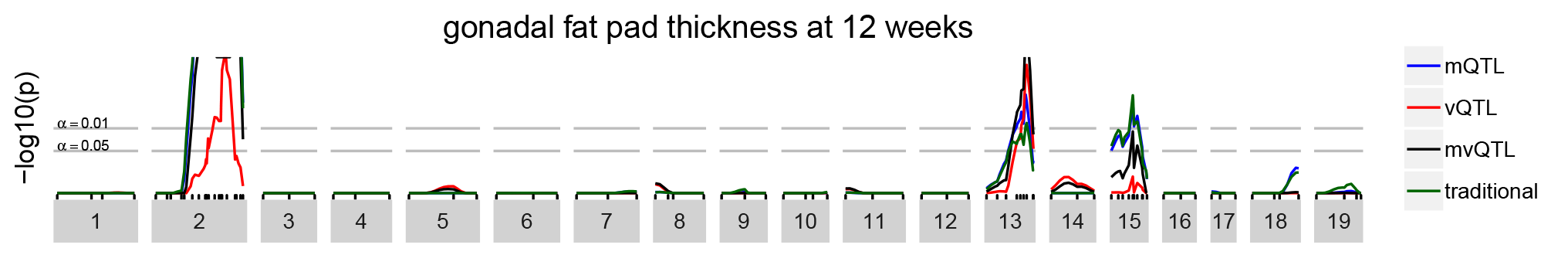

### DGLM Tests for All Phenotypes with BVH

**Figure S14.**
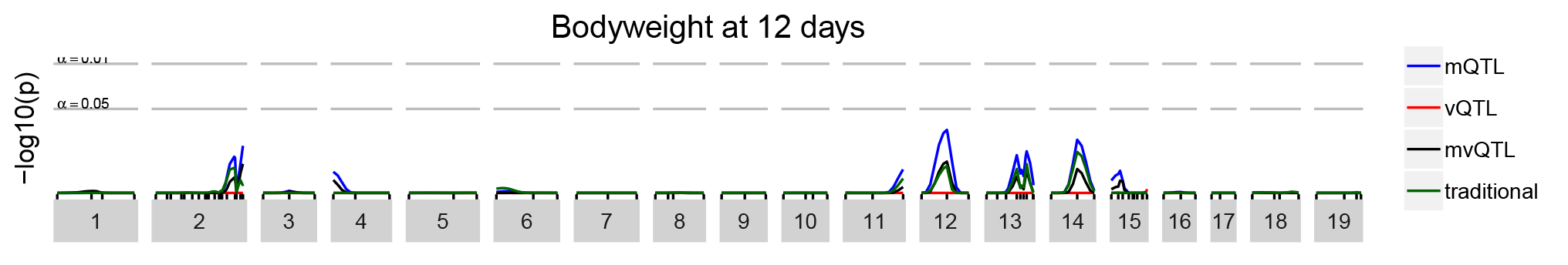
These scans were conducted with the DGLM, accounting for effects of sex and father on variance. For bodyweight at 12 days, no QTL were identified by the SLM or any DGLM-based tests.

**Figure S15.**
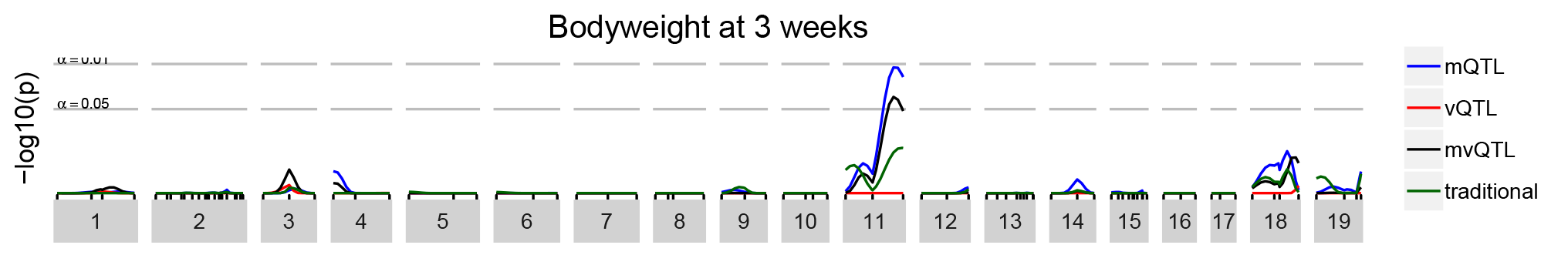
For bodyweight at 3 weeks, an mQTL was identified by DGLM_M_, but not by SLM and no vQTL nor mvQTL were identified. Note that this scan differs slightly from that presented in the **Results** section. This scan accommodates sex as a BVH covariate, as per our “screening” procedure with all phenotypes. Given that the effect of sex on phenotype variance was minimal, we removed it from the analysis in further investigation (but kept the effects of father, which were of much greater magnitude).

**Figure S16.**
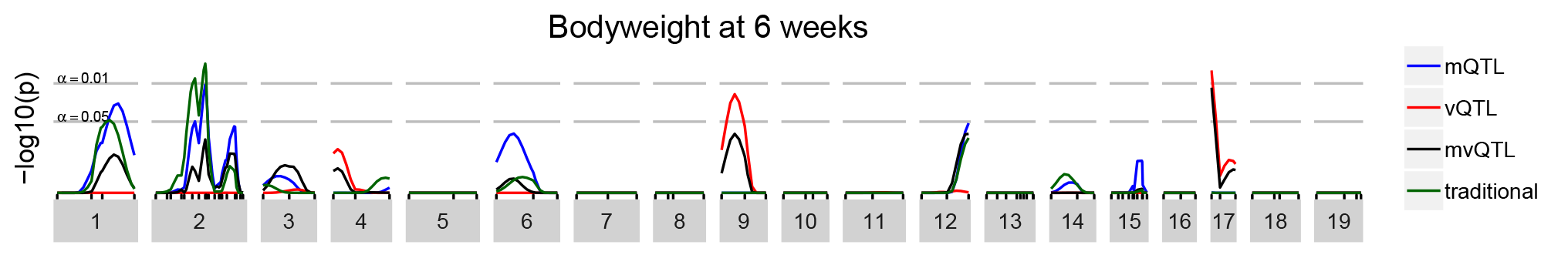
For bodyweight at 6 weeks, SLM identified one mQTL and DGLM_M_ identified the same (chromosome 2). These tests were also concordant in identifying a suggestive mQTL on chromosome 1. DGLM_V_ identified a suggestive vQTL on chromosome 9 and a vQTL on chromosome 17.

**Figure S17.**
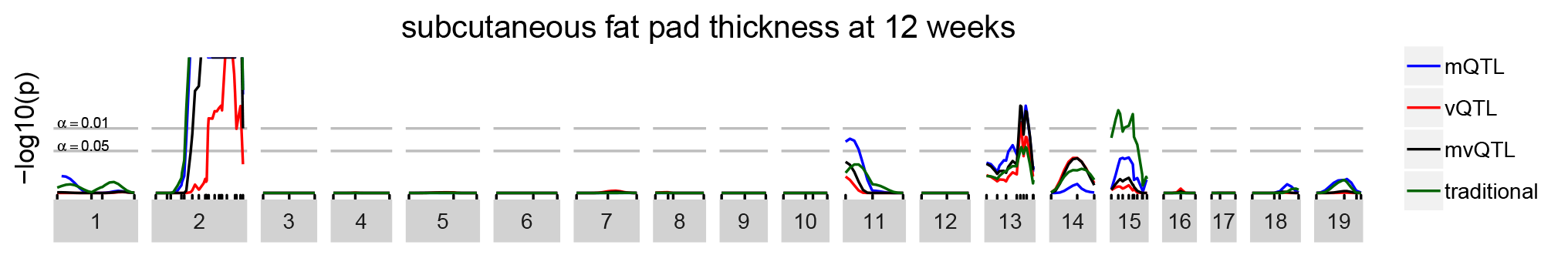
For subcutaneous fat pad thickness at 12 weeks, all tests were highly statistically significant on chromosome 2. On chromosome 13, all tests except SLM were statistically significant. And on chromosome 15, only SLM was statistically significant, reflecting an “un-discovery” due to accommodation of BVH, suggesting that the association identified by SLM was due to a few highly influential observations that were from high-variance sires. Note that the vertical scale is truncated at 10^−4^ for clarity.

**Figure S18.**
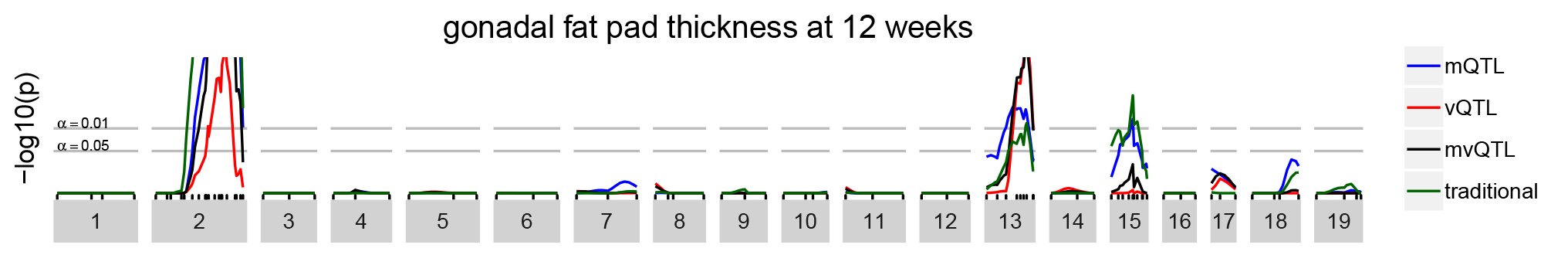
For gonadal fat pad thickness at 12 weeks, all tests were highly statistically significant on chromosome 2. On chromosome 13, all test were statistically significant. An on chromosome 15, both mQTL tests (SLM and DGLM_M_) were statistically significant. Note that the vertical scale is truncated at 10^−4^ for clarity.

